# Differential neural circuit vulnerability to β-amyloid and tau pathologies in novel Alzheimer’s disease mice

**DOI:** 10.1101/2023.04.12.536603

**Authors:** Maria Dolores Capilla-López, Angel Deprada, Yuniesky Andrade-Talavera, Irene Martínez-Gallego, Heriberto Coatl-Cuaya, José Rodríguez-Alvarez, Antonio Rodríguez-Moreno, Arnaldo Parra-Damas, Carlos A. Saura

## Abstract

Alzheimer’s disease (AD) progresses with memory loss and neuropsychiatric symptoms associated with cell specific vulnerability in memory- and emotion-related neural circuits. Neuropathological and synaptic changes are key factors influencing the clinical progression to dementia, but how they cooperate to cause memory and emotional disturbances is largely unknown. Here, we employed pathological, behavioral, expansion microscopy, electrophysiology and transcriptomic approaches to evaluate the effects of amyloid-β (Aβ) and tau on neuropathological progression, synaptic function, and memory and emotional symptoms in amyloid precursor protein (APP), Tau and double novel APP/Tau transgenic mice expressing the mutant human amyloid precursor protein (*APP_Sw,Ind_*) and/or microtubule-associated protein tau (*MAPT*) in excitatory neurons. APP/Tau mice of both sexes show spatial learning and memory deficits associated with synaptic tau accumulation and reduced synaptic proteins and neurotransmission in the hippocampus. By contrast, male and female APP/Tau mice exhibit innate anxious behavior and impaired fear memory extinction linked to Aβ pathology and with absence of synaptic tau in the basolateral amygdala (BLA). Intriguingly, APP/Tau mice show NMDA-dependent long-term potentiation (LTP) deficits in the hippocampus but not in the amygdala. Bulk RNA sequencing reveals region-specific but also common transcriptional changes in response to Aβ/tau pathology, including downregulation of synapse transmission and ion channel activity genes. Importantly, we detected 65 orthologs of human AD risk genes identified in GWAS (e.g., *APOE*, *BIN1*, *CD33*, *CLU*, *PICALM*, *PLCG*2, *PTK2B*, *TREM2*, *SORL1*, *USP6NL*) differentially expressed in the hippocampus and/or BLA of APP/Tau mice, indicating that this APP/Tau model exhibits transcriptional alterations linked to known molecular determinants of AD development. In conclusion, simultaneous development of Aβ and tau neuropathologies in this double APP/Tau transgenic mouse model reproduces synaptic, behavioral, and molecular alterations associated with AD pathophysiology in a region-specific manner. Our findings highlight region-specific pathological effects of Aβ and tau in excitatory neuronal circuits mediating emotional and memory processing, providing evidence that both factors and their molecular cascades should be considered in future AD preventive and therapeutic strategies.

**Graphical abstract:** Age-dependent vulnerability of memory and emotional neural circuits in response to tau and Aβ pathologies.

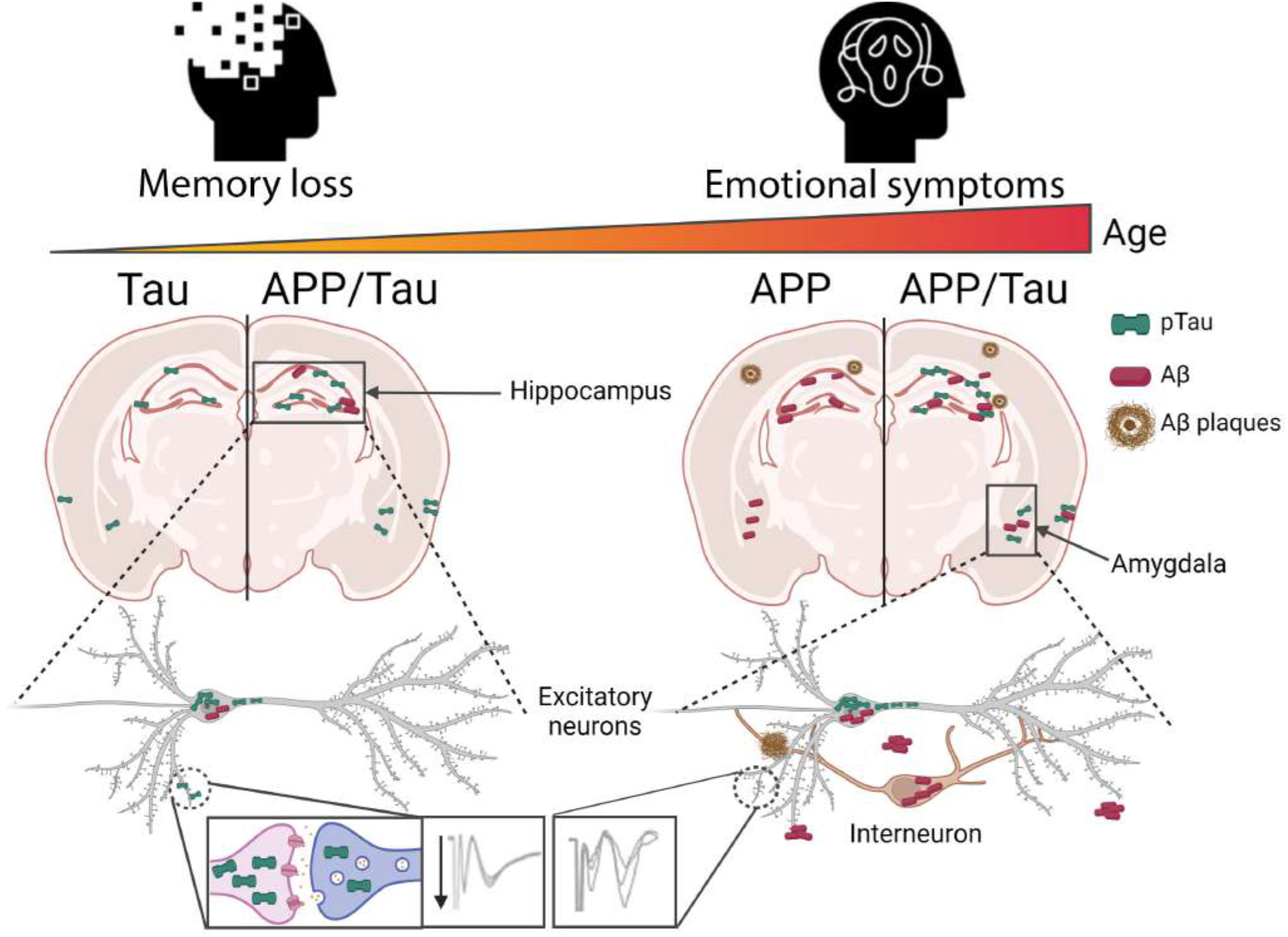

## Introduction

Alzheimer’s disease (AD), the major cause of memory loss in the elderly, is accompanied by neuropsychiatric symptoms, which are observed at the pre-dementia and mild cognitive impairment (MCI) stages and act as early risk factors for conversion to dementia [1, 2]. As part of the medial temporal lobe, the hippocampus and amygdala govern acquisition and expression of episodic and emotional memories, so connectivity changes within these regions underlie cognitive and emotional disturbances at early AD stages [3]. Indeed, the hippocampus and amygdala are activated in response to fear memory [4], but **how these regions cooperate to cause cognitive and emotional changes in AD is unclear**. Among other factors, age and sex directly affect AD clinical and pathological changes across lifespan [5]. Gender differences in cognition and mood disturbances (e.g. depression, anxiety, fear and apathy) are common in AD [6–9], whereas sex-dependent changes in pathology occur during aging [10–13]. In AD mouse models, females are first affected by anxiety and cognitive deficits associated with enhanced neuropathology [14–20]. Compared to males, young 3xTg-AD females show higher anxiety and reduced memory retention [19], whereas sexual dimorphism in behavior, neuropathology and inflammatory molecules are also evident in Tau P301S transgenic mice [20]. **How these pathological factors converge to cause cognitive and neuropsychiatric symptoms is still unclear**, in part because gender and emotional factors are largely underrepresented in basic, pathological and clinical studies [21].

AD is characterized by cerebral accumulation of amyloid-β (Aβ) peptides and microtubule-associated protein tau (MAPT)-containing neurofibrillary tangles (NFTs), and synaptic and neuron degeneration [22]. Recent evidence in humans demonstrates dissociation effects but also a crosstalk of pathological tau- and amyloid-related neural circuit dysconnectivity and memory impairments [23–28]. Several studies demonstrate a crosstalk between tau and amyloid pathologies. Aβ and amyloid plaques increase tau phosphorylation, aggregation and seeding [29–33], and tau inactivation ameliorates Aβ-induced memory deficits independently of amyloid pathology [34, 35]. Importantly, Aβ and tau act synergistically to induce synapse dysfunction and loss [36, 37], a pathological feature that tightly correlates with cognitive decline [38–40]. Pathological tau is present at synapses and induces synapse dysfunction, instability and loss [41–44], and it seems to dominate over Aβ on neural circuit disruption [45, 46]. Aβ and tau affect presynaptic and postsynaptic mechanisms impairing excitatory glutamatergic transmission [41, 47, 48], but whether synaptic tau is responsible and/or synergize with Aβ to induce synapse pathology and behavioral changes is unclear. **Importantly, the crosstalk mechanisms by which Aβ and tau cause synaptic dysfunction and differential vulnerability of emotional and memory neural circuits remain unclear**.

To better elucidate mechanisms and pathological factors contributing to disruption of cognition-and emotion-related neural circuits in AD, we characterized novel double APP/Tau transgenic mice that recapitulate the central neuropathological features of AD. Comparison of single APP and Tau and double APP/Tau transgenic mice revealed that tau and Aβ pathologies preferentially mediate memory loss and emotional disturbances, respectively. Surprisingly, whereas tau exacerbates Aβ-induced LTP deficits in the hippocampus, it counteracts the negative effects of Aβ in LTP in the amygdala of APP/Tau mice. Transcriptomic analyses show differential but also common transcriptional responses related to inflammation and synaptic transmission in the hippocampus and BLA of APP/Tau mice. Simultaneous development of Aβ and tau neuropathologies in this new APP/Tau transgenic mouse model reproduces synaptic, behavioral, and molecular alterations linked to AD pathophysiology.

## Results

### Aβ and tau accumulation in excitatory neurons results in cerebral amyloid and tau pathologies

To investigate the contribution of Aβ and tau on memory and emotional disturbances, we generated and characterized double APP/Tau Tg mice by crossing APP_Sw,Ind_ and Tau P301S Tg mice that express human mutant *APP* and *MAPT*/*tau* transgenes under the excitatory neuron-specific *PDGFβ* and *Prpn* promoters [42, 49]. Accordingly, qRT-PCR analysis of single APP_ RiboTag and Tau_RiboTag mice shows expression of h*APP*, h*Tau* and endogenous *PDGFβ* and *Prpn* translating mRNA transcripts specifically in CaMKIIα excitatory neurons, but not in parvalbumin interneurons (**Supplementary Fig. 1**). Biochemical analysis showed overexpression of human APP (∼2 fold, APP CTF antibody;∼30 fold, 6E10) and tau (∼15-20 fold, D1M9X and TG5 antibodies) and elevated α-CTFs in the hippocampus of 6 month-old APP/Tau mice (**Fig. 1A**; *P* < 0.01). Immunofluorescence analysis revealed high co-localization of human APP/Aβ (6E10) and tau (D1M9X) in excitatory neurons (CaMKIIα) but not inhibitory interneurons (GAD-67 and parvalbumin) of APP/Tau hippocampus, and with a similar staining, except for some colocalization of APP/Aβ with GAD-67, in the BLA of APP/Tau mice (**Figs. 1B**, not shown). At 6 months, APP and APP/Tau mice show APP/Aβ intracellular staining (6E10) in CA1/CA3 hippocampus but not in the dentate gyrus (DG), entorhinal cortex (EC) and BLA (Kruskal-Wallis test, genotype effect, CA1: *P* < 0.01; CA3: *P* < 0.01; DG: *P* < 0.05; EC: *P* < 0.05; BLA: *P* > 0.05) (**Fig. 2A**; **Supplementary Fig. 2**). By contrast, APP/Tau mice show elevated phosphorylated (p)tau (Ser202/Thr205)-positive neurons in hippocampal CA1, CA3 and DG regions, EC and BLA compared with WT, APP and Tau mice (Kruskal-Wallis test, genotype effect, CA1: *P* < 0.05; CA3: *P* < 0.01; DG: *P* < 0.001; EC: *P* < 0.001; BLA: *P* < 0.001; **Fig. 3A**, **Supplementary Fig. 3**). Interestingly, Gallyas silver staining revealed some aggregated tau in EC and BLA neurons only in 6 month-old APP/Tau mice (not shown). These results indicate enhanced Aβ-induced tau pathology in young APP/Tau mice.

**Figure 1.**
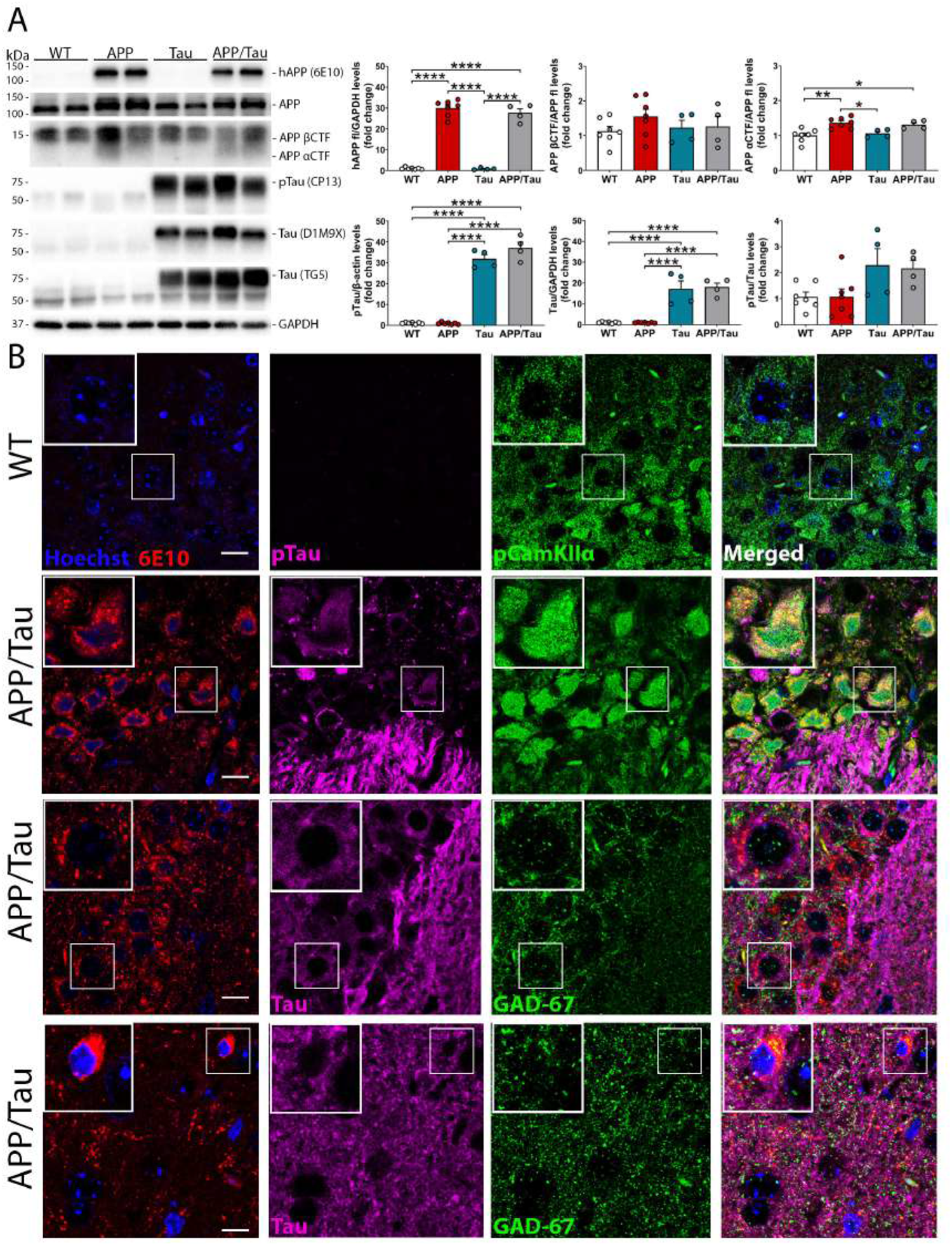
Human APP and tau are expressed in hippocampal excitatory neurons of APP/Tau transgenic mice. **A,** Biochemical analysis and quantification of human APP (6E10 antibody; top), total APP and APP CTFs (CTF antibody) and phosphorylated tau (CP13) and total tau (D1M9X and TG5 antibodies) in the hippocampus of 6 month-old control (WT, white bars), APP (red bars), Tau (blue bars) and APP/Tau (grey bars) mice. Protein levels are normalized to GAPDH or β-actin. Data represent mean ± SEM. n=4-7. Statistical analysis was determined by one-way ANOVA followed by Tukey’s post hoc tests. \**P* < 0.05, \*\**P* < 0.01, \*\*\*\**P* < 0.0001 *vs* the indicated group. **B,** Upper images: Confocal microscope images show colocalization (yellow) of Aβ/APP (6E10; red) and ptau (Ser202; CP13; magenta) in excitatory pyramidal neurons (pCamKII, green) but not inhibitory neuronal projections (GAD67; green) in CA3 hippocampus of 9 month-old APP/Tau mice. Bottom images: Aβ is present in some GAD67-postive cells in the BLA of APP/Tau mice at 9 months as shown in the magnified image. Hoechst staining is depicted in blue. Scale bar: 15 μm.

**Figure 2.**
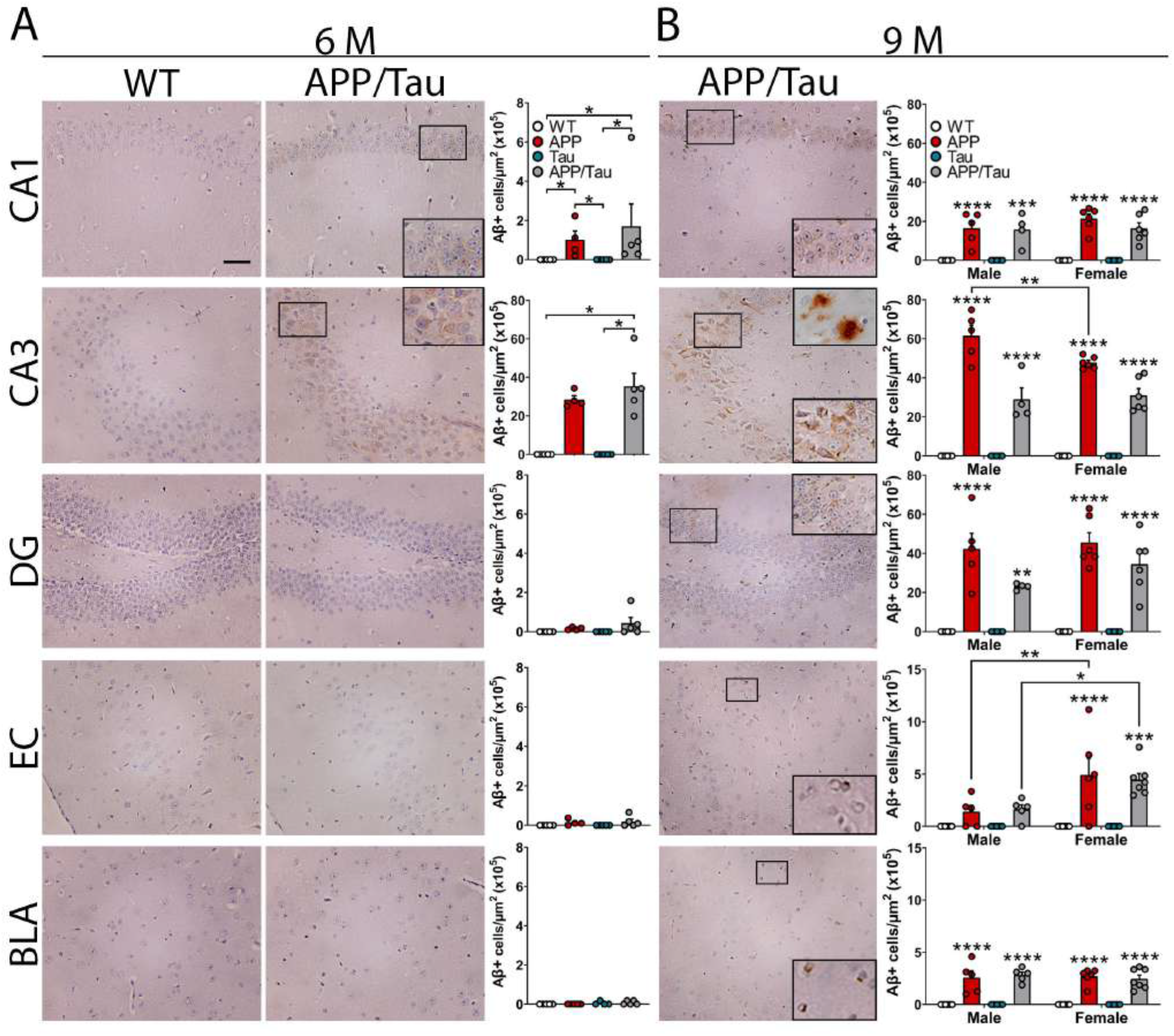
Age-dependent cerebral Aβ pathology in APP/Tau mice. Coronal brain sections of control (WT), APP, Tau and APP/Tau mice at the age of 6 months (**A**) and 9 months (**B**) were stained with anti-human Aβ/APP antibody (6E10). <u>Left images</u>: Representative low and high (insets) magnified images of Aβ-stained neurons in hippocampal CA1, CA3 and dentate gyrus (DG) regions, entorhinal cortex (EC) and basolateral amygdala (BLA). Amyloid plaques in a 9 month-old APP/Tau mouse are visualized in the top right inset of CA3 region. Objective: 20x. Scale bar: 50 μm. <u>Right diagrams</u>: Quantification of Aβ-positive cells in WT (white circles), APP (red circles), Tau (blue circles), and APP/Tau (grey circles) mice. Data represent mean number ± SEM of APP/Aβ positive cells/μm^2^ (x10^5^) (n=3 slices/mouse; 4-7 mice/genotype). Statistical analysis was performed using Kruskal-Wallis test and Dunn’s multiple comparison test (**A**), and two-way ANOVA followed by Sidak’s (sex comparison) and Tukey’s (genotypes comparison within each sex) multiple comparison tests (**B**). \**P* < 0.05, \*\**P* < 0.01, \*\*\**P* < 0.001, \*\*\*\**P* < 0.0001 *vs* WT or the indicated group.

**Figure 3.**
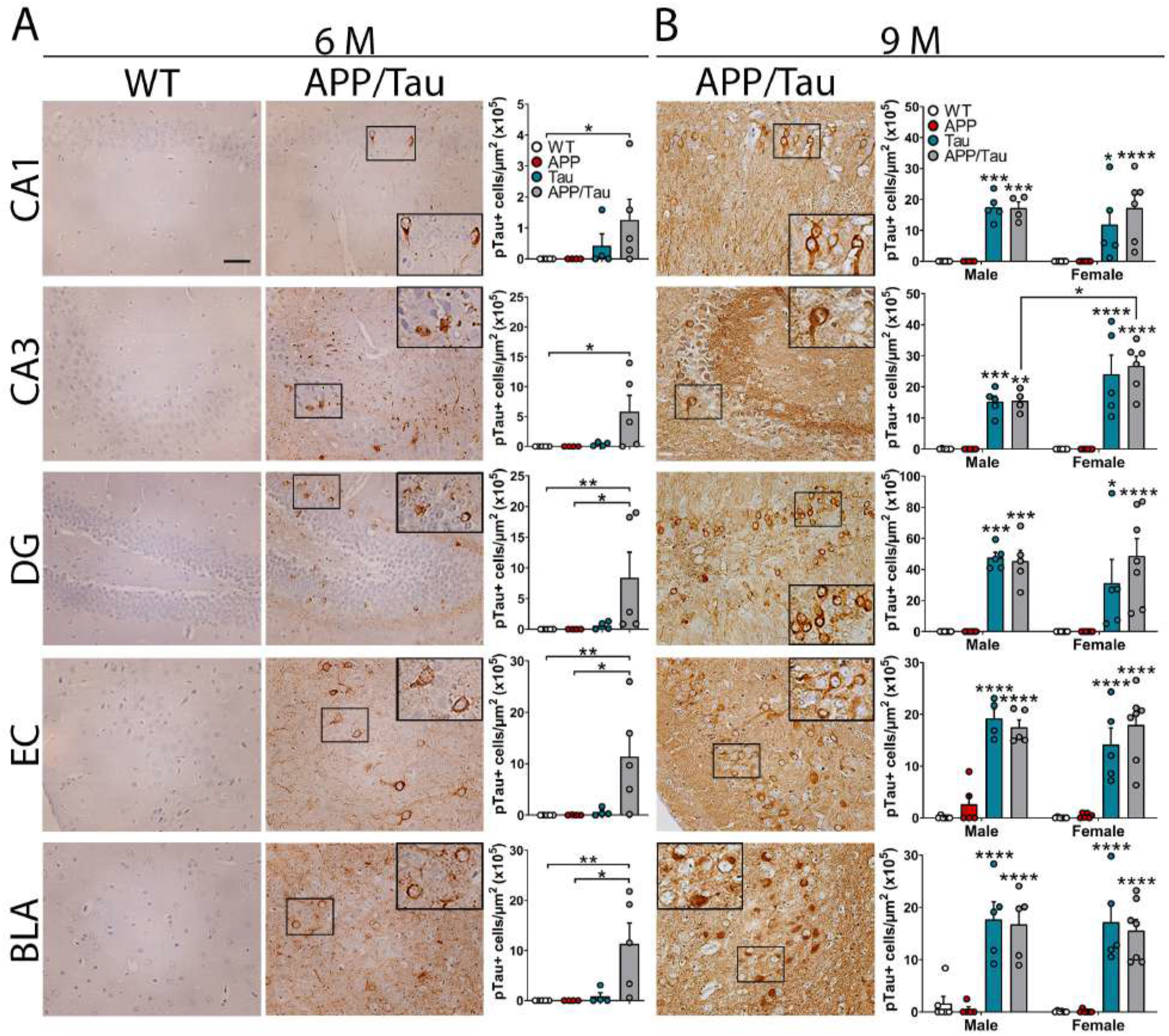
Age-dependent cerebral tau pathology in APP/Tau mice. Brain sections of control (WT), APP, Tau and APP/Tau mice at the age of 6 months (**A**) and 9 months (**B**) were stained with ptau AT-8 or CP13 antibodies. <u>Left images</u>: Representative low and high (insets) magnified images of hippocampal CA1, CA3 and dentate gyrus (DG) regions, entorhinal cortex (EC) and basolateral amygdala (BLA). Objective: 20x. Scale bar: 50 μm. <u>Right diagrams</u>: Quantification of pTau-positive cells in WT (white circles), APP (red circles), Tau (blue circles), and APP/Tau (grey circles) mice. Data represent mean number ± SEM of pTau-positive cells/μm^2^ (x10^5^) (n=3 slices/mouse; 4-7 mice/genotype). Statistical analysis was performed using Kruskal-Wallis test and Dunn’s multiple comparison test (**A**), and two-way ANOVA followed by Sidak’s (sex comparison) and Tukey’s (genotypes comparison within each sex) multiple comparison tests (**B**). \**P* < 0.05, \*\**P* < 0.01, \*\*\**P* < 0.001, \*\*\*\**P* < 0.0001 *vs* WT or the indicated group.

### Tau and Aβ cooperate to induce differential regional synaptic plasticity effects

We next investigated the functional effects of early amyloid and tau pathologies on hippocampal and amygdalar synaptic transmission and plasticity. Input-output curves in Schaffer collateral CA3/CA1 synapses were similarly decreased in all mutant transgenic mice at 6 months (Two-way ANOVA, stimulus effect: F (8,180) = 19.90, *P* < 0.0001; genotype effect: F (3,180) = 34.34, *P* < 0.0001; interaction effect: F (24,180) = 2.24, *P* < 0.01; **Fig. 4A**). Short-term synaptic plasticity (STP), and long-term potentiation (LTP) induction, measured as early-LTP (60 min) and late-LTP (120 min) were significantly impaired in APP/Tau hippocampus compared with the rest of groups (One-way ANOVA, STP: F (3,24) = 4.06, *P* < 0.05; E-LTP: F (3,24) = 3.41, *P* < 0.05; L-LTP: F (3,24) = 7.00, *P* < 0.01; **Figs. 4A-C**). To determine if hippocampal LTP involves pre-and/or post-synaptic mechanisms, we next analyzed paired-pulse facilitation ratio (PPF) showing no significant changes before and after 120 min LTP but baseline PPFs were reduced in APP and APP/Tau mice (F (3,21) = 5.37, *P* < 0.01) (**Fig. 4D**). Whole-cell patch-clamp recordings revealed decreased impaired NMDA responses and NMDA/AMPA ratio in APP/Tau mice (F (3,36) = 3.44, *P* < 0.01; **Fig. 4E**). These results demonstrate that tau and Aβ cooperate to disrupt hippocampal synaptic plasticity through postsynaptic mechanisms.

**Figure 4.**
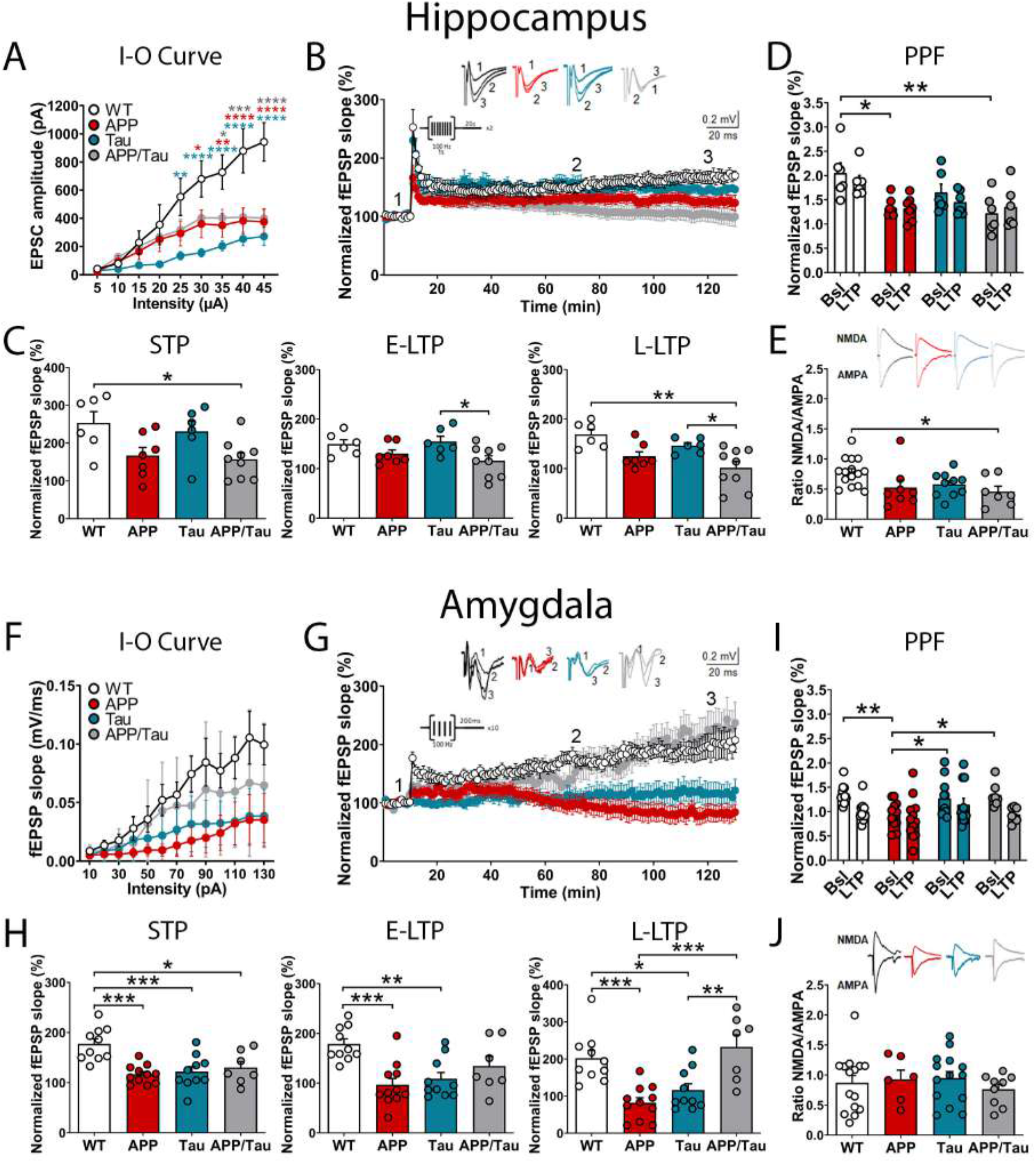
Regional differential effects of tau on Aβ-induced synaptic plasticity impairments. **A,F**, Input-output curves showing ePSC amplitudes (**A**) and fEPSP slope (**F**) *versus* the applied stimulus intensity in CA3/CA1 hippocampal (**A**) and thalamus-lateral amygdala (**F**) synapses of control (WT, white), APP (red), Tau (blue), and APP/Tau (grey) mice. **B,G,** Traces (top) and time course (bottom) of fEPSPs before and after LTP induction in CA3-CA1 hippocampal (**B**) and thalamus-BLA synapses (**G**). **C,H,** Histograms showing short-term synaptic plasticity (STP), early long-term potentiation (E-LTP), late LTP (L-LTP) in CA3-CA1 hippocampal (**C**) and thalamus-BLA synapses (**H**). **D,I,** Paired-pulse facilitation (PPF) does not change after LTP induction in the hippocampus (**D**) or BLA (**I**), and baseline PPFs of APP are decreased in the hippocampus and BLA compared to controls. However, PPF of APP/Tau mice is significantly decreased only in hippocampus. **E,J**, NMDA/AMPA ratio was decreased in hippocampal neurons of APP/Tau mice compared to control, APP, and Tau mice (**E**). In BLA (**J**), NMDA/AMPA ratio was not affected in the transgenic mice. Data represent mean ± SEM of electrophysiological recordings in CA3-CA1 and thalamic-BLA synapses of 9 month-old female control (n = 7-8), APP (n = 5-6), Tau (n = 7-8), and APP/Tau (n = 6) mice. Statistical analysis was determined by one-way (**D,E, I-J**) or two-way (**A,F**) ANOVA followed by Tukey’s post-hoc test. \**P* < 0.05, \*\**P* < 0.01, \*\*\**P* < 0.001, \*\*\*\**P* < 0.0001 *vs* the indicated group. **A-**

In thalamic-lateral amygdala (LA) synapses, basal synaptic transmission was similarly decreased in APP and Tau mice and APP/Tau mice at 6 months (Two-way ANOVA, stimulus effect: F (12, 481) = 3.45, *P* < 0.0001; genotype effect: F (3, 481) = 10.11, *P* < 0.0001; interaction effect: F (36, 481) = 0.3124, *P* > 0.99; **Fig. 4F**). Field recordings revealed decreased STP induction in APP, Tau and APP/Tau mice (F (3,34) = 9.01, *P* < 0.001), whereas deficits in E-LTP and L-LTP were only detected in APP and Tau mice (E-LTP: F (3, 34) = 7.61, *P* < 0.001; L-LTP: F (3, 34) = 11.29, *P* < 0.0001; **Figs. 4G-H**). PPF was unchanged before and after 120 min LTP but baseline PPFs were reduced in APP mice and recovered in APP/Tau mice (F (3,36) = 5.13, *P* < 0.01; **Figs. 4I**). NMDA/AMPA ratios were not significantly affected in the transgenic groups (F (3, 40) = 0.41, *P* > 0.05), although AMPA but not NMDA currents were significantly decreased (*P*<0.001) in APP and Tau mice compared to APP/Tau mice (AMPA, F (3, 40) = 9.34, *P* < 0.0001; NMDA, F (3, 40) = 1.36, *P* > 0.05; **Fig. 4J**). Together, these findings indicate counteracting effects in synaptic plasticity in response to APP/Aβ and tau pathologies in the amygdala of 6 month-old mice.

### Hippocampal-dependent learning and memory deficits in young Tau and APP/Tau mice

Analysis of general behavior in the open field revealed no significant differences among genotypes at 6 months in total travelled distance, crossings, and percentage of time in the center (C) and periphery (P) at days 1 and 2 (∼50% males/females; **Fig. 5A**). This result indicates similar spontaneous locomotor activities and no anxiety-like behavior among groups. Dark/light box and cued fear conditioning (CFC) tests revealed no significant changes in anxiety, neophobia and fear memory among the transgenic lines (Kruskal-Wallis test and two-way ANOVA, *P* > 0.05 (**Figs. 5B, C**). Spatial learning and memory tested in the Morris water maze test showed decreased latencies to reach the hidden platform during training for all groups at 6 months (Two-way ANOVA, training effect: F (4,390) = 37.40, *P* < 0.0001), although Tau and APP/Tau mice exhibited significantly longer latencies starting at day 2 (Genotype effect: F (3,390) = 25.95, *P* < 0.0001; **Fig. 5D**). Swimming speed during training and visible platform performance were similar among all groups (*P* > 0.05), ruling out the possibility of motor and visual deficits. In the probe trial, control mice displayed a preference for the target quadrant (*P* < 0.05), whereas APP, Tau and APP/Tau mice showed reduced target quadrant occupancies (Two-way ANOVA, interaction effect: F (3,156) = 5.86, *P* < 0.001) (**Fig. 5E**). Similar results were found when males and females were analyzed separately (data not shown). In the absence of locomotor and anxiety alterations, hippocampal-dependent spatial learning deficits are associated with tau pathology and synaptic plasticity deficits in APP/Tau mice.

**Figure 5.**
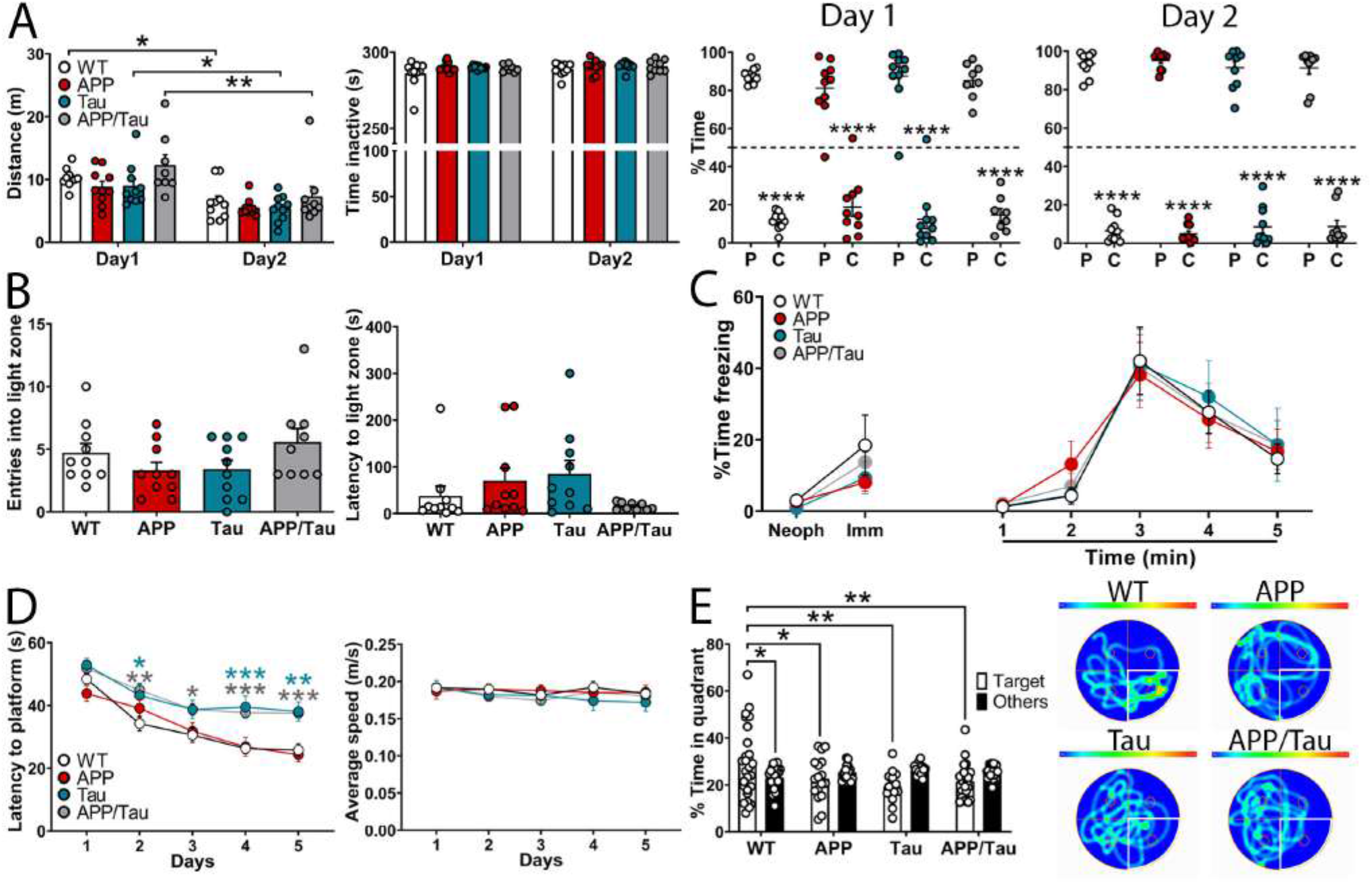
Hippocampal-dependent learning impairments in Tau and APP/Tau mice at 6 months. **A**, General behavior in the open field. Travelled distance (left), time inactive (middle) and percentage of time in the periphery (P) and center (C) of the experimental box (middle and right) during days 1 and 2. Number of mice: control (WT, n= 10; white bars/symbols), APP (n= 10; red bars/symbols), Tau (n= 10; blue bars/symbols) and APP/Tau (n= 9 ; grey bars/symbols) **B,** Number of entries and latencies to light zone in the dark/light box test.. **C**, Associative memory in the fear conditioning test. Freezing responses in the CFC during conditioning (day 1, left) and training (day 2, right) of WT, APP, Tau and APP/Tau mice (n=9–10) at 5-6 months of age. **D,** Spatial memory training in the MWM. Increased escape latencies and unchanged swimming speeds of Tau and APP/Tau mice during spatial training in the MWM. **E**, Day 5 probe trial in the MWM. APP, Tau and APP/Tau mice show disrupted spatial memory retention as revealed by decreased occupancies and trajectories in the target quadrant (marked in white in the heat map). Data represent mean ± SEM. Statistical analysis was determined by Kruskal-Wallis test and Dunn’s multiple comparison test (**B**) or two-way ANOVA (**A,C-E**) followed by Tukey’s post hoc test for genotype comparisons or Sidak’s for day comparison, periphery vs center comparison (**A**) and target comparison (**E**). \**P* < 0.05, \*\**P* < 0.01, \*\*\**P* < 0.001 *vs* the indicated group or periphery (panel A) or control mice (panel D left).

### Age and sex differences in amyloid and tau pathologies in middle-aged APP/Tau mice

To consider the possible effects of transgenes and gender on AD-related pathological and behavioral changes during aging, we next analyzed separate cohorts of middle-aged (9 months) mice. APP and APP/Tau mice of both sexes show similar intracellular Aβ and amyloid plaques CA1, DG, EC and BLA, except for decreased Aβ-positive neurons in the CA3 layer of APP/Tau mice (**Fig. 2B**, **Supplementary Fig. 5)**. This result indicates that, compared to tau, Aβ pathology starts first in the hippocampus and then in the EC and BLA. Remarkably, Aβ-positive cells in EC were significantly increased in APP and APP/Tau females compared to males (*P* < 0.05). Besides accumulation of extracellular Aβ, Aβ/APP staining was also localized in some GAD-67-positive interneurons in BLA (**Fig. 1B**, bottom). Compared with young mice, phosphorylated tau (Ser202, CP13) was prominently increased in neuronal cell bodies and fibers of hippocampus and cortex of both Tau and APP/Tau mice (∼20-40 fold change) and, particularly in CA3 region of APP/Tau females (Two-way ANOVA, CA3, genotype effect: F (3,35) = 43.57, *P* < 0.0001; Sex effect: F (1,35) = 7.87, *P* < 0.01) (**Fig. 3B, Supplementary Fig. 6**). These results demonstrate age, gender and regional differences in amyloid and tau pathologies in APP/Tau mice. Notably, whereas tau appears simultaneously in distinct brain regions, Aβ pathology appears in the hippocampus and then in the EC and BLA of APP/Tau mice.

### Age-dependent emotional disturbances in APP and APP/Tau mice

Evaluation of general behavior in the open field revealed no significant differences in travelled distances among genotypes in male and female mice at 9 months, indicating similar ambulatory locomotor activities (**Figs. 6A,D**). However, APP/Tau females showed more activity and increased time in the periphery on day 1 (Two-way ANOVA, zone effect: F (1,58)= 1818, *P* < 0.0001; interaction effect: F (3, 58)= 10.30, *P* < 0.0001), and APP males showed increased time in the periphery at day 2 (Zone effect F (1,72)= 1571, *P* < 0.0001; interaction effect: F (3, 72)= 5.138, *P* < 0.01) (**Figs. 6A, D**). In the dark/light box test, 9 month-old APP and APP/Tau mice of both sexes, but not Tau mice, showed reduced entries (One-way ANOVA, male: F (3,36) = 6.88, *P* < 0.001; females: F (3,29) = 5.16, *P* < 0.01), and higher latencies (except male APP/Tau mice; male: F (3,36) = 2.98, *P* < 0.05; females: F (3,29) = 4.47, *P* < 0.05) into the light compartment (**Figs. 6B, E**), suggesting increased anxiety in male and female APP and APP/Tau mice. In CFC, innate freezing responses before shock (neophobia) and immediately after shock are similar in female groups (Two-way ANOVA, genotype effect: *P* > 0.05), whereas males show a main significant effect of genotype (F (3,108) = 5.23, *P* < 0.01) and treatment (F (2,108) = 15.70, *P* < 0.0001) (**Figs. 6C, F**). Notably, male and female APP mice show increased freezing behavior after foot shock, suggesting increased immediate response to the aversive stimulus. At 24 h, all groups showed similar and significant CS-induced freezing responses indicating fear memory consolidation. However, there was a significant effect of genotype and tone during extinction training (males: genotype effect: F (3,324) = 9.57, *P* < 0.0001; tone effect: F (8,324) = 9.51, *P* < 0.0001; females: genotype effect: F (3,306) = 12.56, *P* < 0.0001; tone effect: F (8,306) = 11.48, *P* < 0.0001) (**Figs. 6C, F**). Particularly, APP and APP/Tau mice of both sexes show significant enhanced average of freezing responses compared to control and Tau mice (genotype effect, male: F (3,24) = 7.00, *P* < 0.01; females: F (3,24) = 12.51, *P* < 0.0001) (**Figs. 6C,F**). These results indicate that APP and APP/Tau mice develop emotional symptoms associated with Aβ accumulation in the amygdala.

**Figure 6.**
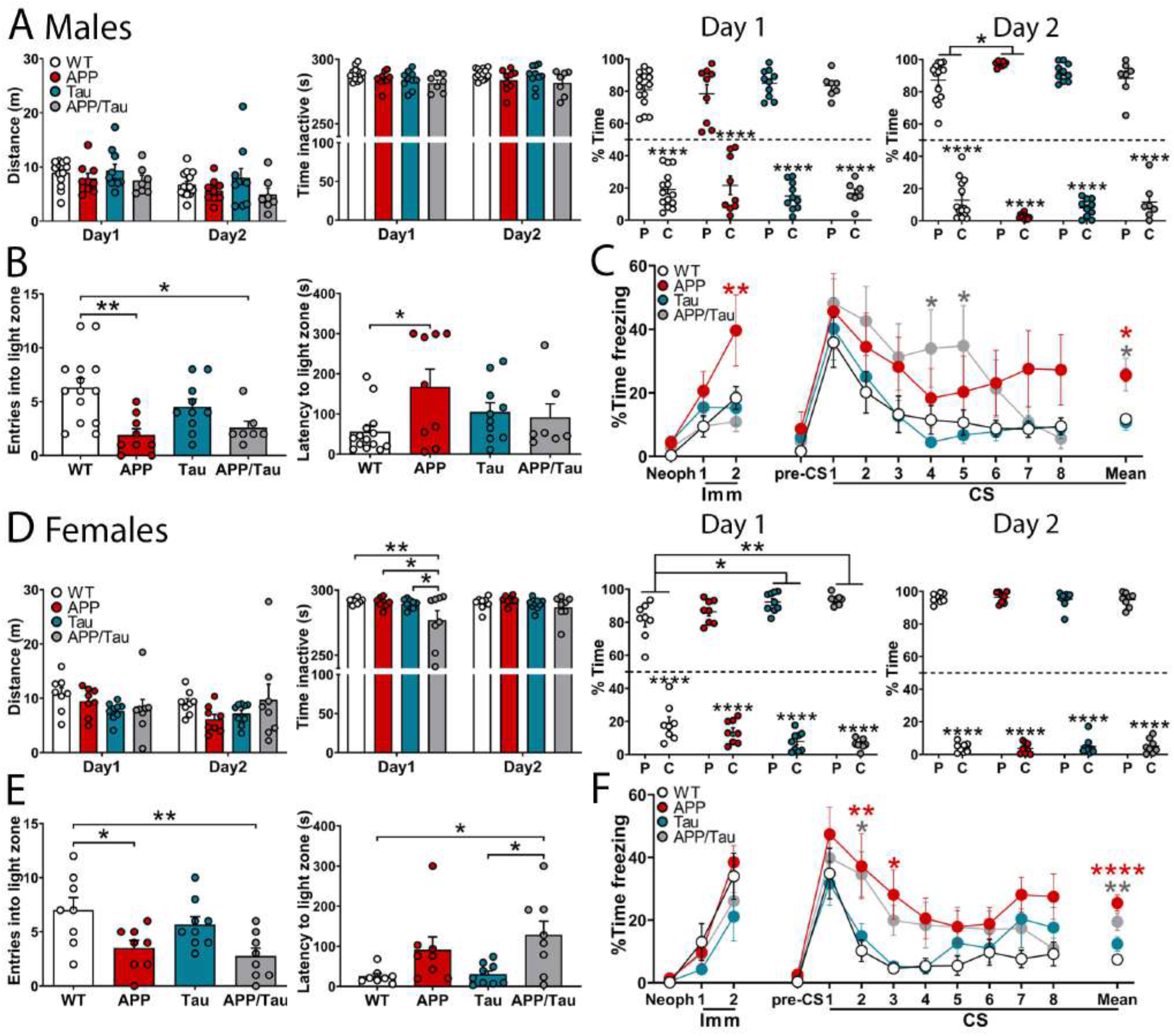
Emotional disturbances in APP and APP/Tau mice at 9 months. **A, D,** Behavior of male (**A**) and female (**D**) mice in the open field. Travelled distance (left), time inactive (middle) and percentage of time in the periphery (P) and center (C) of the box (right) in the experimental groups during days 1 and 2. **B, E,** Anxiety behavior of male (**B**) and female (**E**) mice in the dark/light test. APP and APP/Tau mice show decreased number of entries and/or increased latencies to the light zone in the dark/light box test. **C, F**, Associative memories of male (**C**) and female (**F**) mice in the CFC test. Freezing responses during conditioning (day 1) and training (CS 1: conditioned stimulus 1) or extinction (CS 2-8) on day 2 in the CFC. The mean ± SEM of time freezing (%) during CS (period from 2 to 8) is represented at the right. In all cases, data represent mean ± SEM. Number of mice: control (WT, n= 14 male, n= 8 female; white bars/symbols), APP (n= 9 male, n= 8 female; red bars/symbols), Tau (n= 10 male, n= 9 female; blue bars/symbols) and APP/Tau (n= 7 male, n= 8 female; grey bars/symbols). Statistical analysis was determined by one-way (**B, E, mean of CS 2-8 of C, F**) or two-way (**A, C, D, F**) ANOVA followed by Tukey’s post hoc test for genotype comparisons or Sidak’s for day comparison and periphery vs center comparison (**A**). \**P* < 0.05, \*\**P* < 0.01, \*\*\*\**P* < 0.0001 *vs* the indicated group, the periphery (panel **A**) or control mice (panels **C, F**).

### Common transcriptional responses in the hippocampus and BLA of APP/Tau mice

To identify region-specific transcriptional responses and biological pathways affected by Aβ/tau pathology, we next performed bulk RNA sequencing (RNAseq) in the hippocampus and BLA of female WT and APP/Tau mice at 9 months. Differential expression analysis of gene expression revealed significant changes in 5,985 (46 % down, 54 % up) and 2,168 (42% down, 58% up) genes in the hippocampus and BLA, respectively, of APP/Tau mice (**Fig. 7A**). GO analysis revealed that upregulated differentially expressed genes (DEGs) genes in the hippocampus (e.g., *Cd14, Cd44, Cd68, Lamp1/2, Pycard*) were associated with immune responses (neutrophil and cytokine terms) whereas downregulated genes (e.g., *Cacna1a/b/c, Camk2a*, *Gria1/2/3*, *Grik1/2, Grin1*, *Grin2a/b*, *Kcnc1/3/4, Ptk2b, Ryr1*) were linked to axon development and neurotransmission (glutamate receptor and cation channel activity). In BLA, the upregulated genes (e.g., *Apoe, Arhgap9*, *Cd14, Cd33, Cst3, Ctsd, Ctss, Clu, Ilr1, Itgam, Irak1, Stat3*) were involved in inflammatory responses (neutrophil and cytokine terms), Aβ and metabolism, and the downregulated genes (e.g., *Camk2a*, *Dlgap1/4, Gria2/3*, *Grin1, Homer1, Lrrk2, Mapk8ip2, Mink1, Shank1, Syn1*) were related to glutamate neurotransmission and GTPase regulation (**Fig. 7B**). A high number of DEGs (1,423: 24% hippocampus, 65% BLA) were coordinately altered in both regions in APP/Tau mice. Importantly, we detected 65 DEGs that are mouse orthologs of human AD risk genes, identified in GWAS studies (e.g., *APOE*, *BIN1*, *CD33*, *CLU*, *MS4A4A*, *PICALM*, *PLCG*2, *PTK2B, SLC24A4, SORL1*, *TREM2*, *USP6NL*) [50, 51], altered in the hippocampus and/or BLA of APP/Tau mice (**Fig. 7B; Supplementary Table 1**). These results suggest shared and region-specific transcriptional responses to Aβ/tau pathology related to inflammation and synapse function in the hippocampus and BLA of APP/Tau mice.

**Figure 7.**
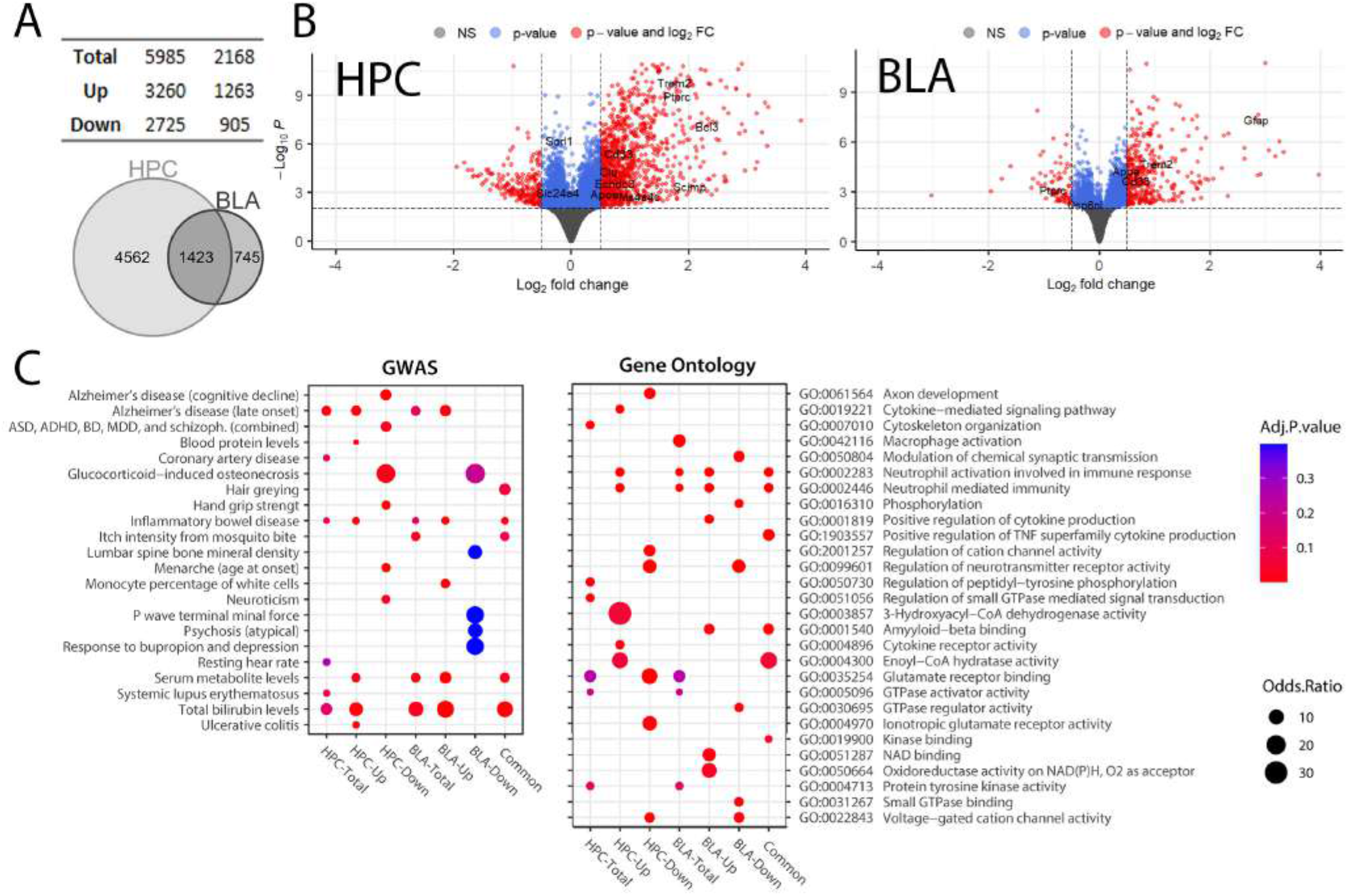
Bulk RNAseq reveals differential and common gene expression signatures associated with Alzheimer’s disease in hippocampus and BLA of APP/Tau mice. **A,** Number and Venn diagram illustrating the total, up and down differentially expressed transcripts in the hippocampus and BLA of 9 month-old female APP/Tau mice compared to controls (WT). **B,** Volcano plots showing the fold change of genes differentially expressed in the hippocampus and BLA of APP/Tau mice (threshold of < 0.05 adjusted *P* value, abs log_2_FC >0.1). Some significant deregulated AD risk genes are indicated (see also **Supplementary Table 1**). **C**, **D**, Functional enrichment of differentially expressed genes in APP/Tau hippocampus and BLA. The plots describe the six first significant functionally enriched pathways from Genome-wide association studies (GWAS) catalog (2019) (left) and the three first significant functionally enriched pathways from GO Biological Process (2021) and GO Molecular Function (2021) (right) databases. Human orthologs were used in the analysis with GWAS catalog database.

### Synaptic tau is associated with synapse pathology in the hippocampus of APP/Tau mice

Synapse dysfunction and synapse-to-nucleus signaling contribute to cognitive and emotional impairments in neurodegenerative and neuropsychiatric diseases [52]. To examine whether changes in synaptic transmission and plasticity, and synaptic gene networks are related to altered synaptic proteins, we next performed biochemical and tissue expansion microscopy analyses. Biochemical analyses revealed increased tau in total and synaptic fractions but no changes of synaptic proteins in hippocampal lysates of 6 month-old APP/Tau mice (**Fig. 8A**). Notably, presynaptic proteins (synaptophysin, syntaxin 1A) and glutamatergic postsynaptic proteins (CRTC1, GluN1, Homer1, PSD95) were decreased in purified hippocampal synaptosomes and/or pre- and post-synaptic fractions of APP/Tau mice (**Fig. 8A**). Immunofluorescence staining in expanded brain sections revealed colocalization of ptau (Ser 202) with the postsynaptic protein Homer1 in the hippocampus but not in the BLA of APP/Tau mice at 6 months (**Fig. 8B; Supplementary Fig. 6)**. These changes in synaptic proteins that occur in parallel with pre- and post-synaptic tau accumulation suggest that synaptic pathological tau contributes to hippocampal synapse pathology.

**Figure 8.**
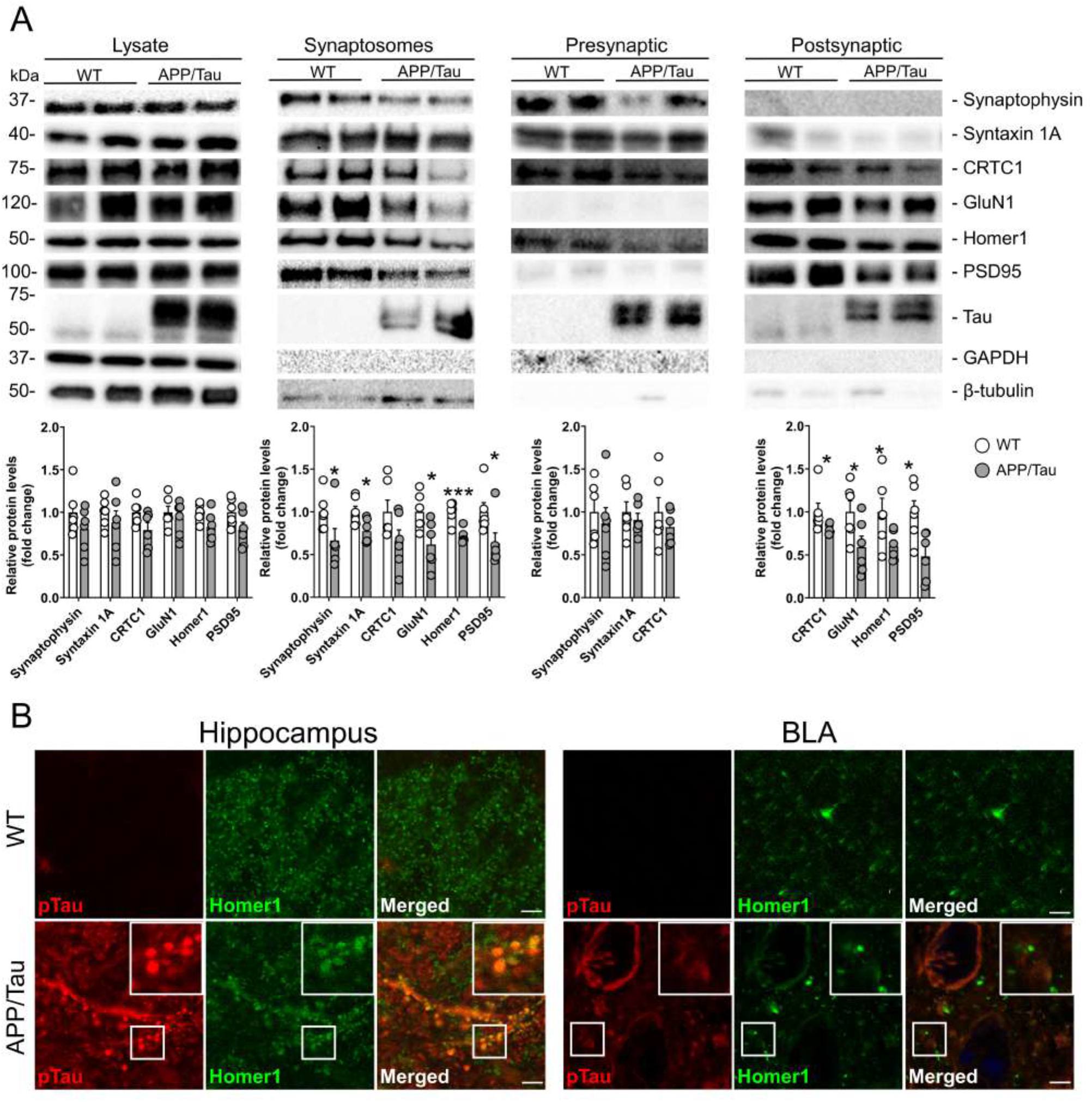
Accumulation of synaptic tau and decreased synaptic proteins in the hippocampus of APP/Tau mice. **A**, Biochemical analysis of tau and synaptic proteins in total hippocampal lysates, and purified synaptosomes and pre- and post-synaptic fractions from WT and APP/Tau mice at 6 months (n=6 mice/group). Synaptic proteins are unchanged in total APP/Tau hippocampal lysates, whereas pre- and post-synaptic proteins are decreased mainly in purified synaptosomes and post-synaptic fractions of APP/Tau mice coinciding with increased synaptic tau accumulation (D1M9X antibody). Data represent mean fold change ± SEM. Values were normalized to GAPDH or β-tubulin (total lysate) or total protein (synaptosomes, pre- and post-synaptic fractions). Statistical analysis was determined by two-tailed Student’s t-test or two-tailed Mann-Whitney test according to the normality test. \**P* < 0.05, \*\**P* < 0.01, \*\*\**P* < 0.001. **B**, Confocal microscope images of expanded hippocampal and BLA sections (expansion factor 3.63x) of 6 month-old WT and APP/Tau mice showing colocalization of postsynaptic Homer1 (green) and p-tau (Ser 202; red) in CA1 hippocampus. Scale bars: 10 μm and 36.30 μm (expanded).

## Discussion

Human neuroimaging studies demonstrate that Aβ and tau impair synapse function and cause axonal damage either of which may lead to disruption of neuronal connectivity and cognitive deficits in AD [23, 25–27]. Our extensive pathological, behavioral, electrophysiological and transcriptomic characterization of novel APP/Tau mice demonstrates that Aβ and tau in excitatory neurons disrupt synapse function and integrity leading to differential vulnerability of memory- and emotion-related circuits. Cognitive and emotional disturbances are temporally associated, respectively, with accumulation of tau in the hippocampus and Aβ in the amygdala of APP/Tau mice, suggesting age- and region-specific dissociation of memory and emotional symptoms in AD. Remarkably, whereas Aβ and tau cooperate to disrupt synaptic transmission and plasticity in the hippocampus, they counteract each other to maintain synaptic function and plasticity in the BLA. Accordingly, transcriptional profiling indicates regional differences but also coordinated gene and cellular networks related to inflammation and synapse function in response to Aβ and tau pathological changes. These results indicate differential regional vulnerability in cellular pathways mediating memory and emotional processing in response to Aβ and tau pathologies (**see Graphical Abstract**).

Our experimental approach allows precise dissection of regional and gender effects of Aβ and tau pathologies during aging. Considering that age and gender play modifying roles on cognitive decline, elucidating sex differences is critical for designing strategies to detect, prevent and develop personalized AD treatments. APP /Tau mice of both sexes develop similar cerebral pathologies, but compared to males, aged females show elevated Aβ and tau pathologies specifically in EC and CA3 hippocampus, respectively. This result is relevant considering that life expectancy and elevated amyloid and/or tau pathologies are associated with higher prevalence of dementia in women [21], and emotional disturbances associated with verbal cognitive decline occur more often in middle-aged females than in males [53]. Interestingly, whereas phosphorylated tau appears simultaneously in distinct brain regions, Aβ pathology starts in the hippocampus and extends later to the EC and BLA, coinciding with innate anxious behavior and impaired fear memory extinction. This reinforces the deleterious effect of Aβ on amygdala-associated emotional disturbances [54, 55], and contrasts with the mild AD pathological changes observed in humanized APP, tau or Aβ *knockin* mice [56–58]. In agreement with previous studies [30–32, 59–61], Aβ potentiates tau pathology in the hippocampus, EC and BLA in young APP/Tau mice, whereas mutant tau does not apparently affect APP processing and amyloid pathology [29, 34, 35]. This result is relevant considering that amyloid pathology is critical for tau-mediated hippocampal and amygdala dysfunction and memory deficits in non-demented subjects [23]. It is plausible, however, that tau affects amyloid pathology-associated dystrophic neurites or neurodegeneration [62, 63], and could antagonize the effects of Aβ on hyperexcitation causing aberrant neuronal activity [46, 64]. The underlying mechanisms by which Aβ affects tau pathology are still unclear, but neuronal activity, microglia, and genetic variants could be critical players [65–67].

The amygdala and hippocampus cooperate on long-term memories while playing independent roles in emotion and declarative memory processing [68]. It is interesting that impaired spatial and fear emotional memories are temporally linked to accumulation of pathological tau and Aβ in the hippocampus and amygdala, respectively. This may explain why anxiety and cognitive deficits are tightly associated with both pathologies in 3xTg-AD, APP and tau transgenic mice [14–16, 19, 54, 55]. Surprisingly, middle-aged APP/Tau male mice do not show such as dramatic innate anxious behavior as APP mice (**Fig. 6C**). Although the neurobiological basis of this difference is unclear, it is possible that, as occurs in the cortex, tau mediates suppression of Aβ-induced hyperactivity in the amygdala [46]. These emotional alterations are associated with the presence of Aβ in BLA excitatory and inhibitory interneurons, pointing to a role of Aβ-induced excitatory/inhibitory imbalance in amygdala-dependent emotional impairments. This is intriguing considering that Aβ and/or phosphorylated tau in BLA alter amygdala function in AD and FTD [54, 69], and that neuropsychiatric symptoms are tightly associated with tau in dementia during aging [70, 71]. Considering that anxiety is reversed by anti-tau therapeutic treatments [72–74], the apparent lack of tau effects on anxiety and fear extinction in APP/Tau mice could be explained by dominating effects of Aβ over tau and/or differences in cell-specific transgene expression, genetic background and/or age of the transgenic lines.

Tau pathology in memory-related circuits tightly predicts memory decline in older adults [25, 75], and accumulation of tau at synapses contributes to synaptic plasticity deficits in Tau P301S mice [36, 37, 42, 47, 76]. Accordingly, Aβ and tau act synergistically to impair hippocampal synaptic transmission and plasticity in APP/Tau mice. This could involve a postsynaptic mechanism as indicated by reduced NMDA/AMPA ratio, although the possibility for a presynaptic effect of tau, as suggested by PPF decrease, cannot be rule out [42, 76, 77]. A link between synaptic tau and synapse pathology is reinforced by the presence of tau at hippocampal synapses of APP/Tau mice coinciding with impaired synaptic plasticity and reduced presynaptic and postsynaptic proteins, including the glutamate receptor-related proteins CRTC1, GluN1, Homer1, PSD95. Based on the synergistic effect of Aβ and tau on synaptic dysfunction [78], it is plausible that Aβ could promote tau-mediated hippocampal synapse pathology. Alternatively, intracellular Aβ may cause synapse dysfunction independently of APP and tau overexpression as observed in humanized Aβ mice [57]. It is intriguing, however, that the presence of both pathologies counteracts synaptic transmission and plasticity deficits in the amygdala of single transgenic mice. One possibility is that the pathology is not fully established in the amygdala at 6 months, as suggested by the absence of synaptic tau in BLA, with the subsequent lack of effect on anxiety. When the pathology progresses (e.g., at 9 months), deregulation of the glutamatergic circuitry emerges as in the hippocampus of APP knock-in mice [79]. Additionally, the possibility of increased excitability accompanying the response after the LTP protocol in APP/Tau mice should not be excluded. Interestingly, our transcriptome profiling revealed altered expression of genes related to synaptic function and neurotransmission in both regions (**Fig. 7**), although ultimately synaptic plasticity deficits were compensated in the amygdala of double APP/Tau mice. Indeed, synaptic genes are tightly associated with AD clinical status and disease progression, being upregulated in mild cognitive impairment in multiple brain regions, including the hippocampus and prefrontal cortex, and then decline as the disease progresses [80–82]. Interestingly, single-cell transcriptomics recently revealed deregulation of synaptic transmission and signaling pathways in excitatory and inhibitory neurons, astrocytes and oligodendrocytes in AD brain ([83], for a review). The fact that we detected 65 DEGs orthologs of human AD risk genes identified in GWAS studies in APP/Tau mice strongly supports that our mouse model exhibits transcriptional alterations linked to known molecular determinants of AD development. Integrative co-expression-based analysis of murine and human bulk and single-cell transcriptomics datasets will be useful to discern relevant cell-type specific genes in specific neural circuits affected by AD pathology.

## Materials and methods

### Mice

Littermate non-transgenic control (WT), APP, Tau and APP/Tau Tg mice were obtained by crossing heterozygous APP_Sw,Ind_ (line J9; C57BL/6) and Tau P301S (line PS19, JAX #008169; C57BL/6) mice expressing the familial AD-linked K670N/M671L and V717F *APP* and frontotemporal dementia (FTD)-linked P301S *Tau* mutations in neurons under the *Platelet-derived growth factor β-chain* (*PDGFβ*) and *prion protein* (*Prpn*) promoters, respectively, respectively [42, 49]. We used separate 6- and 9-10 month-old male and female groups. WT, APP and Tau mice expressing either CaMKIIα^Cre^;RiboTag or PValb (PV)^Cre^;RiboTag were used for gene profiling in excitatory- or parvalbumin (PValb)-positive neurons (unpublished). RiboTag RPL22 Tg mice (C57BL/6) express hemagglutinin (HA)-tagged ribosome-bound mRNAs specifically in Cre-recombinase-expressing cells [84]. Mice were grouped in ventilated cages in standard housing conditions (12 h light/12 h dark cycle). Procedures to reduce the number, pain, suffering, and distress of experimental animals are approved by the Animal and Human Ethical Committee of the Universitat Autònoma de Barcelona (CEEAH/DMAH: 2895/10571 and 4750/10839) following European Union regulations (2010/63/EU).

### Behavioral tests

Mice were kept in the original cages and randomly allocated into experimental groups. Anxiety-like behaviors and exploratory activity were studied in the open-field (OF) and dark/light box test [85, 86]. 6 month-old mice were tested in cued fear conditioning (CFC) in a novel conditioning chamber (15.9 x 14 x 12.7 cm) equipped with a bright light (1850 Lux) (context A) (Med Associates, St. Albans, Vermont) for 3 min (neophobia) before the onset of a cue tone (conditioned stimulus, CS, 2800 Hz and 80 dB; 30 sec) that ended with a footshock (unconditioned stimulus, US: 0.8 mA, 2 sec). Mice remained 2 min in the chamber (immediate freezing), and freezing behavior was examined at 24 h in a novel chamber (context B) before (pre-CS; 2 min) and during (CS; 3 min) tone presentation. An independent middle-aged group (9-10 months) was conditioned in context A for 3 min before 2x pairings of the CS with a US (1 mA, 2 sec). Freezing behavior was examined automatically (Video Freeze Software, Med Associates) in the 2 min inter-pairing interval (immediate freezing 1) and after footshock (2 min, immediate freezing 2) [54]. 24 h later, and after 3 min (pre-CS), extinction training was conducted using 16 x CS presentations (30 sec each, 5-sec inter-CS) in context B [87]. Morris water maze (MWM; pool: 1.2 m diameter) during five days (6 trials/day; 60 sec/trial) followed by a 24 h probe trial (60 sec) was analyzed using ANY-maze software [54]. Experimenters were blind to the genotypes of the mice.

### Biochemical analysis and subcellular fractionation

For biochemical analysis, half hippocampus was lysed in cold-lysis buffer (62.5 mM Tris hydrochloride pH 6.8, 10 % glycerol, 5 % β-mercaptoethanol, 2.3 % sodium dodecyl sulfate (SDS), 5 mM NaF, 100 μM Na_3_VO_4_, 1 mM EDTA, 1 mM ethylene glycol tetraacetic acid (EGTA), 1 μM okadaic acid and 50 μM cyclosporin A) containing protease and phosphatase inhibitors and boiled [55, 88]. Sucrose gradient fractionation of synaptosomes and pre- and post-synaptic fractions of individual hippocampi was performed as described [89]. Proteins were resolved on SDS-polyacrylamide gel electrophoresis (PAGE) and detected by immunoblotting with the following antibodies: APP C-terminal fragment (CTF; Sigma, A8717), human Aβ (6E10) (BioLegend, SIG-39320), synaptophysin (Sigma, S5768), β-tubulin (Sigma, T9026), and GAPDH (Life Technologies, AM4300), APP CTF (Sigma, A8718), tau (Cell Signaling, D1M9X; TG5), pTau (Ser202, CP13), Homer1 (Synaptic Systems, 160003), PSD95 (Cell Signaling, 2507), GluN1 (Merck-Millipore, MAB363), Syntaxin 1A (Santa Cruz sc-12736), and CRTC1 (Cell Signaling #2501). Chemiluminescent bands were captured and quantified in a linear range using a ChemiDoc MP System and ImageLab 5.2.1 software (Bio-Rad).

### Cell-type and bulk transcriptional profiling

Hippocampi of APP or Tau CaMKIIα^Cre^;RiboTag or PValb^Cre^;RiboTag mice (6 months) were lysed in lysis buffer (50mM Tris pH 7.5, 100mM KCl, 12mM MgCl_2_, 1% Nonidet P-40, protease inhibitor, 1mM DTT, 1mg/ml heparin, 100ug/ml cycloheximide, 200U/ml RNAse out). Polyribosomes were immunoprecipitated (IP) with Anti-HA.11 Epitope Tag antibody (BioLegend) plus protein A/G magnetic beads, and total RNAs from input and IP were extracted using Invitrogen PureLink RNA mini kit (Thermo Fisher Scientific). Purified RNA was reverse-transcribed and amplified by qRT-PCR. IP/Input ratio of human *APP* and *Tau*, and murine *PDGF*β and *Prpn* genes was calculated by the ΔCt method.

Bulk RNA-seq transcriptional profiling was performed in 9 month-old females (n=4-6/group) using dissected hippocampi and basolateral amygdala (BLA; 1 mm Ø micropunch) from eight and two consecutive frozen coronal brain slices (300 μm). Total RNA was extracted using the RNeasy Micro Kit (Qiagen) and analyzed using Agilent 2100 Bioanalyzer. cDNA libraries were prepared from 50 ng (hippocampi) or 10 ng (amygdalae) RNA using the NEBNext® Ultra™ II RNA Library Prep Kits for Illumina (New England Biolabs). Libraries were sequenced (40M bp sequencing depth, paired-end 2 x 50 bp) on an Illumina NextSeq 2000 instrument. Adapter removal and trimming were performed with Trimmomatic (v0.39) and RNA-seq data analyses, including alignment to the reference genome (mm39) and differential expression analysis, were performed using QuasR/Rhisat2 and DESeq2 packages in Bioconductor (v3.16)[90, 91]. Gene ontology (GO) and functional enrichment analyses were performed using enrichR package in Bioconductor [92].

### Electrophysiological recordings

Coronal brain slices (350 μm) containing the hippocampus or the two amygdalae of 6 month-old female mice were maintained (30-34 °C) in cerebrospinal fluid (CSF) consisting of (in mM): 126 NaCl, 3 KCl, 1.25 KH_2_PO4, 2 MgSO4, 2 CaCl_2_, 26 NaHCO_3_, and 10 glucose (pH 7.2, 300 mOsmL^-1^) [93]. Field excitatory postsynaptic potentials (fEPSPs) were recorded in CA1 hippocampus or lateral amygdala after stimulation (0.2 Hz) of the Schaffer collateral or the thalamic pathway, respectively [93, 94]. Basal synaptic transmission was examined by stimulus-response curves (input-output) by applying 5-10 μA increasing stimulation. In paired-pulse experiments, two consecutive stimuli separated by 40 msec were applied. Data were filtered at 3 kHz and acquired at 10 kHz. Synaptic plasticity was induced by applying a theta burst stimulation (TBS) of 10 bursts of four pulses each (100 Hz, 100 ms duration, 200 ms inter-burst interval; amygdala) or two stimulus trains (100 pulses 100 Hz /train, 20 sec interstimulus; hippocampus). After stable baseline (10 min) TBS was delivered and short-term plasticity and early-(60 min) and late-(120 min) long-term potentiation (LTP) were analyzed.

Whole-cell patch-clamp recordings in the soma of visually identified CA1 pyramidal cells or amygdalar excitatory neurons using IR-DIC microscopy were recorded with a patch-clamp amplifier (Multiclamp 700B). Patch electrodes were pulled from borosilicate glass pipette and had a resistance of 4–7 MΩ when filled with an intracellular solution (pH 7.2–7.3, 290 mOsm L^-1^) containing (in mM): cesium chloride, 120; HEPES, 10; NaCl, 8; MgCl_2_, 1; CaCl_2_, 0.2; EGTA, 2 and QX-314. AMPAR- and NMDAR-EPSCs were evoked at 0.2 Hz and recorded in voltage-clamp mode holding the membrane potential at 70 and +40 mV, respectively. All signals were low-pass filtered at 2 kHz, acquired at 10 kHz, digitized, and stored using Digidata 1322A and pCLAMP 10.4 software (Molecular Devices, CA, USA).

### Immunohistochemistry

Brains were fixed in 4% phosphate-buffered paraformaldehyde (2h) before paraffin embedding. Deparaffinized coronal brain sections (5 μm) were antigen retrieved with citrate buffer (10 mM, pH 6.0) for tau, formic acid (60%, 5 min) for Aβ or both for tau and Aβ. Sections were incubated with monoclonal anti-phosphorylated (p)tau (Ser202, CP13, 1:50; Ser202/Thr205, AT8, ThermoFisher MN1020, 1:50) or anti-Aβ (6E10, 1:1000) antibodies and biotin-conjugated anti-mouse secondary antibody (1:200) before applying the DAB peroxidase substrate kit (Vector laboratories) and imaging with a Nikon Eclipse 80i microscope. For triple immunostaining, deparaffinized sections were incubated with Aβ antibody (6E10, 1:1000), which was blocked (not shown) using a biotin-conjugated Fab anti-mouse antibody (Jackson Immuno Research), before incubation with: 1) mouse pTau (CP13, 1:50) and rabbit anti-pCamKIIα (Abcam, Ab 32678; 1:500) antibodies, or 2) rabbit Tau (D1M9X; 1:600) and mouse GAD-67 (Abcam, Ab 26116; 1:200) antibodies. Sections were incubated with Cy3-conjugated streptavidin (1:1000) and either AlexaFluor-488/647-conjugated goat anti-mouse or anti-rabbit IgGs (1:400) and Hoechst (1:10,000; Invitrogen). For tissue expansion, frozen brain sections (100 μm) labelled with anti-pTau (CP13, 1:50), rabbit anti-Homer 1 (1:300) and/or Bassoon (Abcam, Ab82958; 1:300) antibodies and Hoechst, were incubated overnight with Acryloyl-X SE (Thermofisher, A20770) and then with gelling solution (2h, 37°C) as reported [95]. Sections were digested with proteinase K and cleared and expanded with deionized water washes. Confocal images were obtained with a Zeiss Axio Examiner D1 LSM700 laser scanning microscope (Carl Zeiss Microcopy) and analyzed with ImageJ 6.0.

### Statistical analysis

Statistical analysis (Prism software, GraphPad) was performed using parametric one- or two-way Analysis of Variance (ANOVA) or non-parametric Kruskal-Wallis tests according to D’Agostino-Pearson omnibus normality test. For multiple comparisons, we used Sidak’s or Tukey’s post hocs for parametric tests and Dunn’s post hoc for non-parametric test. Parametric (unpaired two-tailed Student’s t-test) or nonparametric (unpaired two-tailed Mann-Whitney) tests were used when only two groups were compared. *P* values less than 0.05 were considered significant.

## Significance

Discerning specific effects and interactions of Aβ and tau in the human brain is complex and unreliable, therefore, AD mouse models are valuable preclinical tools to investigate sex and age effects on pathological and functional changes in vulnerable neural circuits. In this study, we characterized a preclinical APP/Tau mouse model suitable to investigate cell vulnerability and pathological changes in response to Aβ and tau in excitatory neural circuits. Pathological, behavior, electrophysiological, expansion microscopy and transcriptional analyses pinpoint region-specific pathological roles of Aβ and tau in excitatory neuronal circuits underlying emotional and memory processing, respectively. Transcriptional profiling reveals coordinated shared and region-specific transcriptional responses, including 65 AD risk genes, in the hippocampus and BLA of APP/Tau mice, indicating that this APP/Tau model exhibits region-specific transcriptional changes linked to known molecular determinants of AD development. This study has important implications for translational research on pathological mechanisms underlying differential effects of Aβ and tau on vulnerable memory- and emotion-related neural circuits in AD.

## Acknowledgements

We thank Dr. E. Sanz and A. Quintana for providing the RiboTag mice, and M Puigdellívol for critical reading of the manuscript. We thank the Servei d’Estabulari, Servei de Genòmica i Bioinformàtica and INc Histology and Microscope facilities of UAB for technical support.

## Authors contributions

MDCL, APD and CAS designed and coordinated the study. MDCL designed and performed the pathological, immunofluorescence and behavioral analyses. MDCL, AD and APD performed the transcriptomics, synapse biochemical characterization and expansion microscopy. YAT, IMG, HCC and ARM designed and performed the electrophysiological experiments. MDCL, AD, APD, ARM, JRA and CAS discussed and interpreted the data. MDCL, APD and CAS designed and wrote the manuscript with input of all authors. The authors read and approved the final manuscript.

## Availability of data and materials

Data and materials, including specific experimental protocol information, are available under request. Raw transcriptomics data will be available once the study is published.

## Declarations

### Ethics approval

All animal procedures conducted in the framework of this project were performed according to the European Union regulations (86/609/EEC, 2010/63/EU) in accordance with guidelines and protocols approved by the Animal and Human Ethical Committee (CEEAH) of the Universitat Autònoma de Barcelona and local government of Catalunya (CEEAH/DMAH: 2895/10571 and 4750/10839). Procedures were conducted to reduce the number, pain, suffering, and distress of experimental animals.

### Competing ineterests

The authors declare no financial and non-financial competing conflicts of interest in relation to this work.

