## Supplemental Figures for "Differential neural circuit vulnerability to β-amyloid and tau pathologies in novel Alzheimer’s disease mice"

**Supplementary Figures**

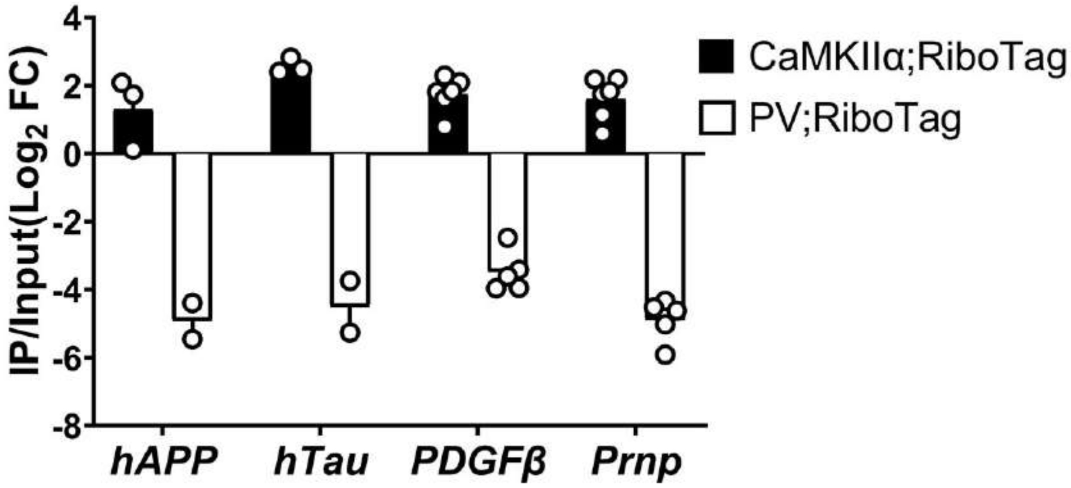


**Supplementary Figure 1. Human *APP* and *MAPT*/*Tau* transgenes are expressed in glutamatergic neurons.**

qRT-PCR analysis showing expression of *PDGFβ* and h*APP* and *Prpn* and h*MAPT*/*Tau* in glutamatergic neurons (CaMKIIα+; black bars) but not parvalbumin interneurons (PV+; white bars) in the hippocampus of APP or Tau CaMKIIα$^{Cre}$ RiboTag or PValb$^{Cre}$ RiboTag mice, respectively. $Log_2$ fold change (FC)/input values indicate enriched (positive values) or lack of expression (negative values) in the specific cell type. Data represent mean ± SEM.

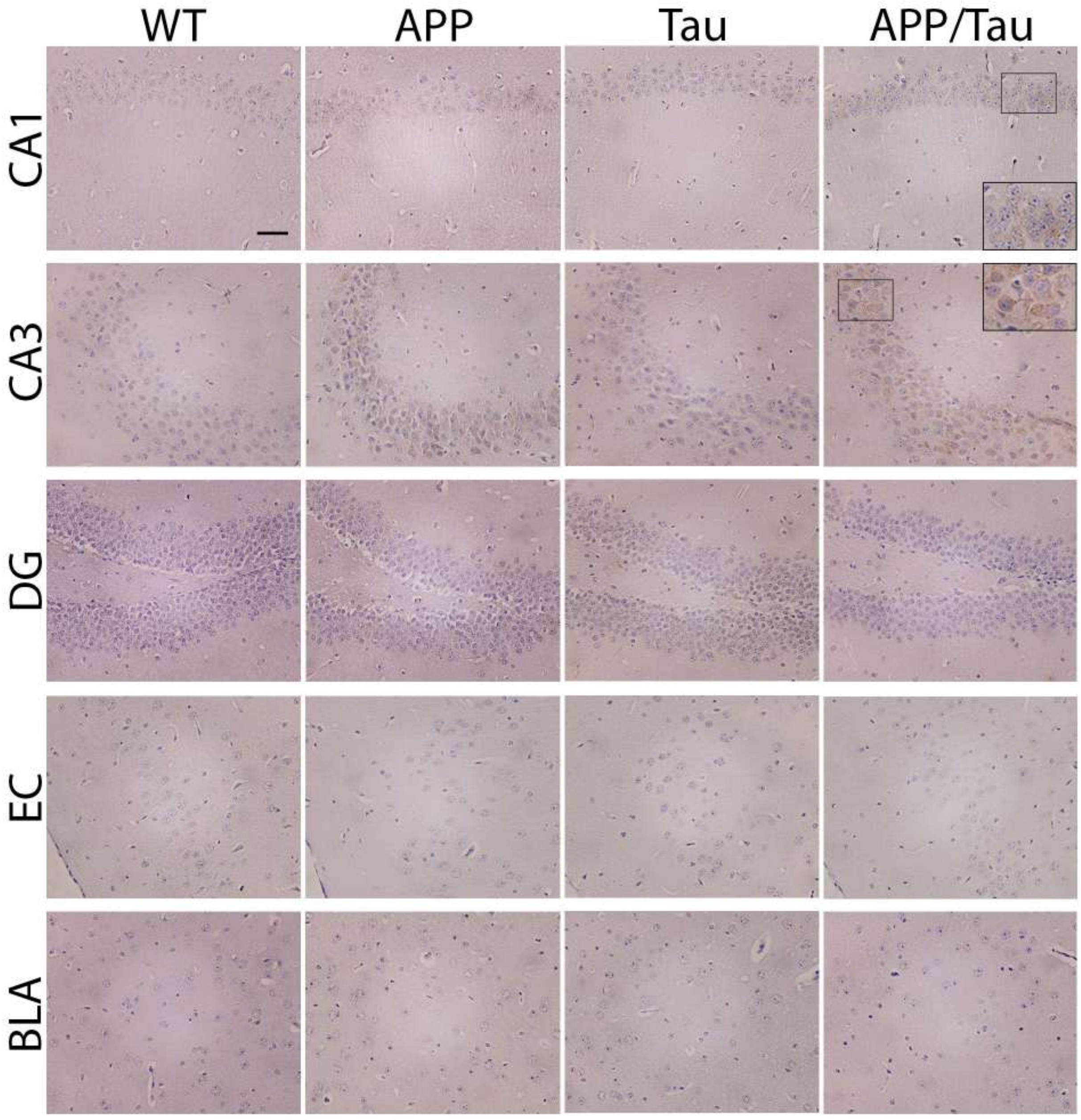


**Supplementary Figure 2. Age-dependent cerebral Aβ pathology in APP and APP/Tau mice at 6 months.**

Coronal brain sections of control (WT), APP, Tau and APP/Tau mice at the age of 6 months were stained with anti-human Aβ/APP antibody (6E10). Representative low and high (insets) magnified images of Aβ-stained neurons in CA1, CA3 and dentate gyrus (DG) hippocampal regions, entorhinal cortex (EC) and basolateral amygdala (BLA) are shown. Objective: 20x. Scale bar: 50 μm

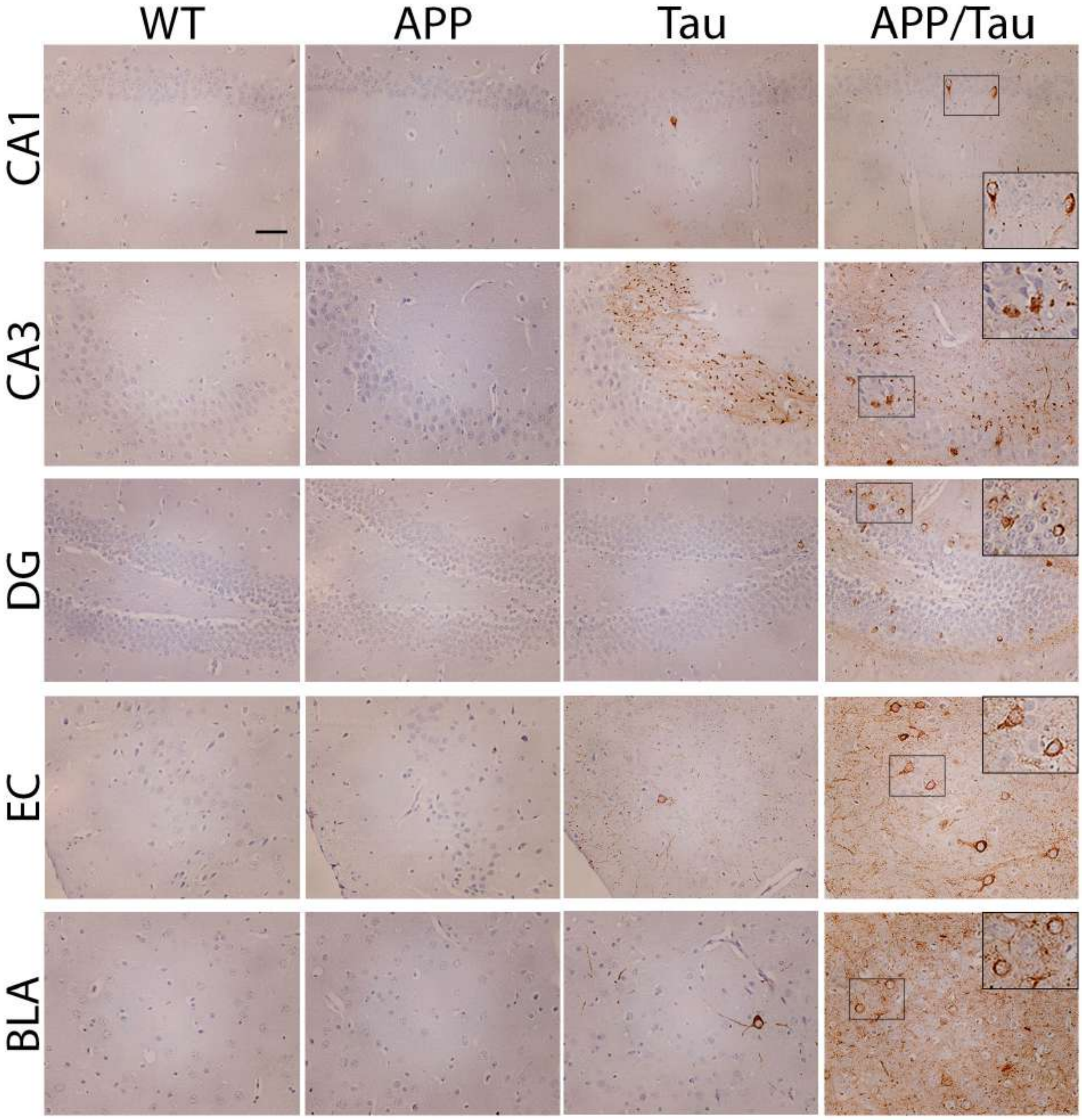


**Supplementary Figure 3. Age-dependent cerebral tau pathology in Tau and APP/Tau mice at 6 months.**

Coronal brain sections of control (WT), APP, Tau and APP/Tau mice at the age of 6 months were stained with anti-phosphorylated tau AT-8 antibody (Ser202/Thr205). Representative low and high (insets) magnified images of tau-stained neurons in CA1, CA3 and dentate gyrus (DG) hippocampal regions, entorhinal cortex (EC) and basolateral amygdala (BLA) are shown. Objective: 20x. Scale bar: 50 µm

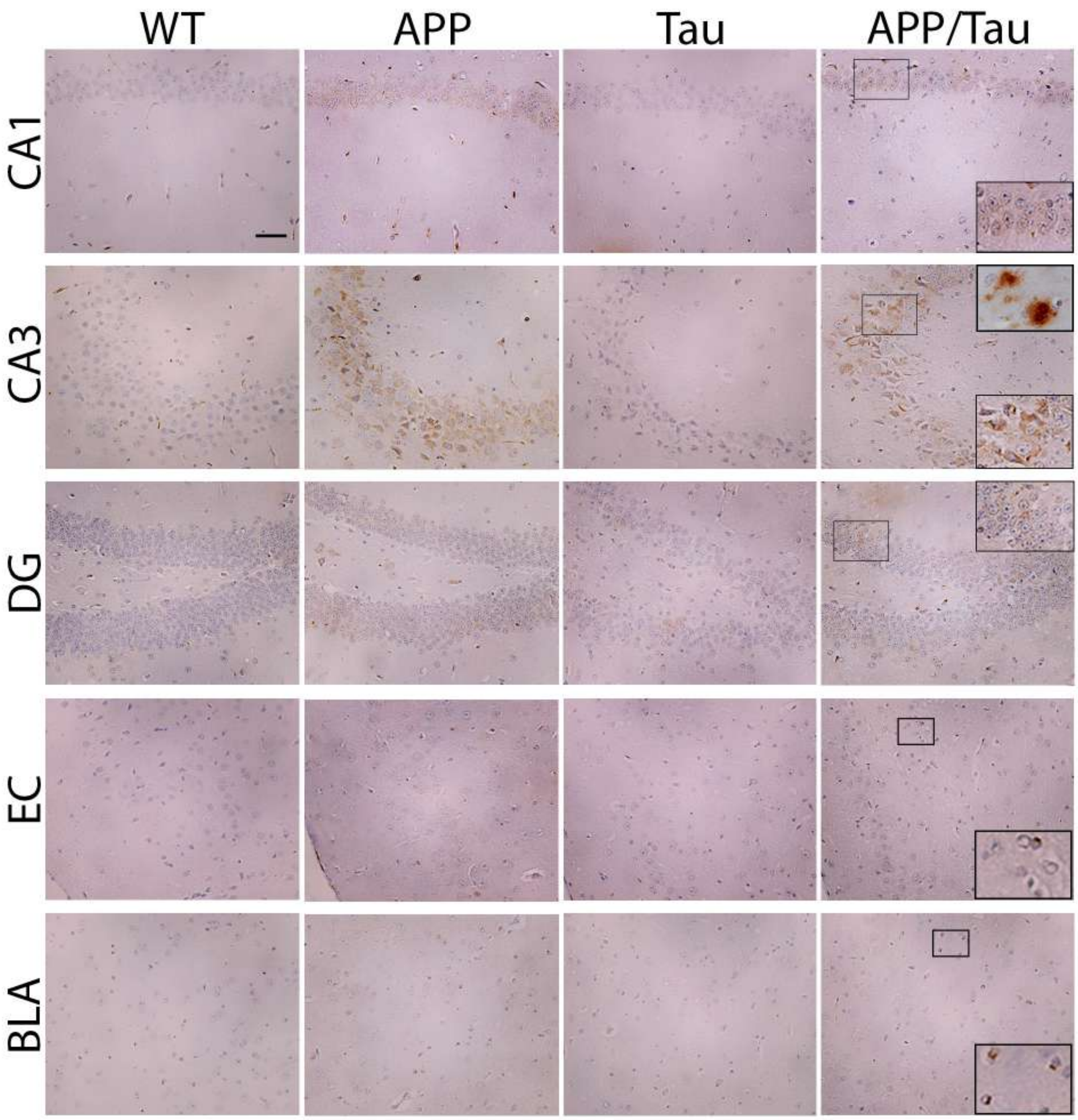


**Supplementary Figure 4. Age-dependent cerebral Aβ pathology in APP and APP/Tau mice at 9 months.**

Coronal brain sections of control (WT), APP, Tau and APP/Tau mice at the age of 9 months were stained with anti-human Aβ/APP antibody (6E10). Representative low and high (insets) magnified images of Aβ or amyloid plaques in CA1, CA3 and dentate gyrus (DG) hippocampal regions, entorhinal cortex (EC) and basolateral amygdala (BLA) are shown. Objective: 20x. Scale bar: 50 μm

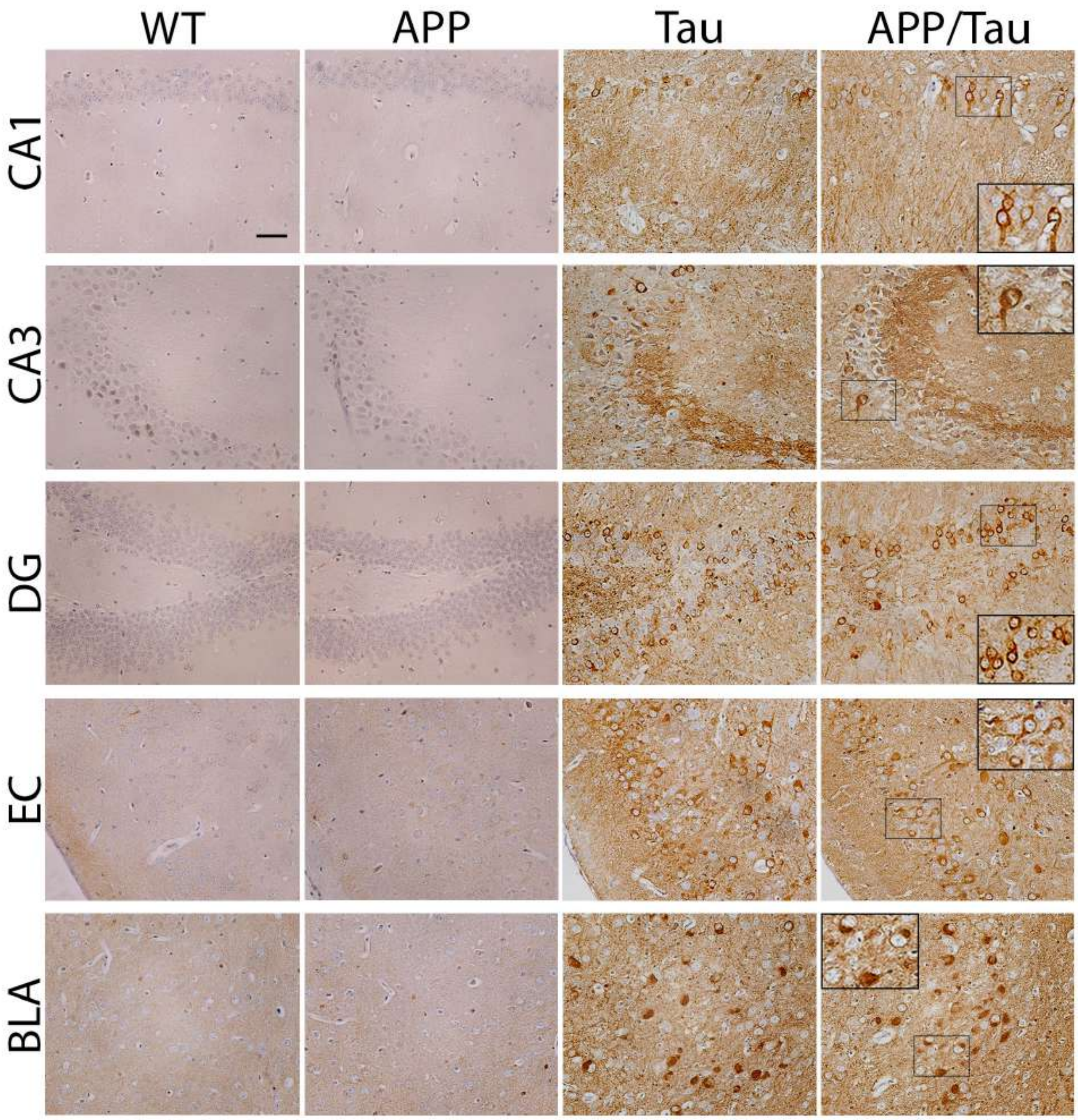


**Supplementary Figure 5. Age-dependent cerebral tau pathology in Tau and APP/Tau mice at 9 months.**

Coronal brain sections of control (WT), APP, Tau and APP/Tau mice at the age of 9 months were stained with anti-phosphorylated tau CP13 antibody (Ser202). Representative low and high (insets) magnified images of tau-stained neurons in CA1, CA3 and dentate gyrus (DG) hippocampal regions, entorhinal cortex (EC) and basolateral amygdala (BLA) are shown. Objective: 20x. Scale bar: 50 μm

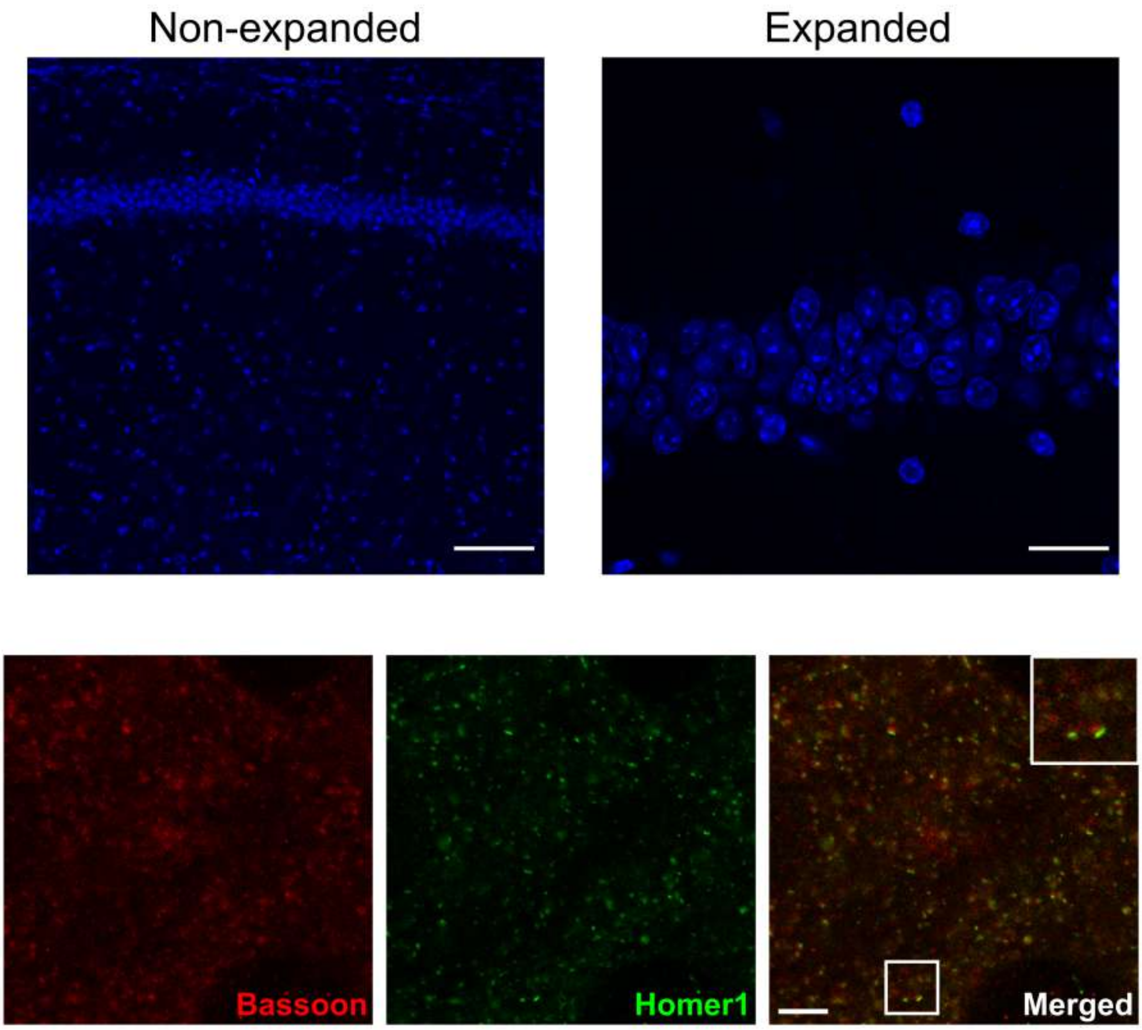


**Supplementary Figure 6. Physical magnification of brain samples using expansion microscopy.**

Confocal fluorescence images showing nuclei (Hoechst dye, blue) of pre- (left) and post-expanded (right) CA1 hippocampal mouse sections (Top), and Bassoon (red) and Homer1 (green) staining in an expanded brain section. Scale bars: Top: pre- versus post-expanded images: 100 µm (physical size post-expansion, 363 µm). Bottom: 10 µm.

**Supplementary Table 1. Deregulated AD risk genes in the hippocampus and BLA of APP/Tau mice**

| | |
|---|---|
| *ADAMTS20* | *NCS1* |
| *AHNAK* | *NKAIN2* |
| *AOX1* | *PDE7B* |
| *APOC1* | *PICALM* |
| *APOE* | *PLCG2* |
| *ARHGAP20* | *PPP1R37* |
| *ATXN7L1* | *PRRC2C* |
| *BCAS3* | *PTK2B* |
| *BCL3* | *RHBDF1* |
| *BIN1* | *RIN3* |
| *CD2AP* | *SCIMP* |
| *CD33* | *SEC24B* |
| *CDON* | *SLC24A4* |
| *CELF1* | *SLC4A8* |
| *CLU* | *SORL1* |
| *COBL* | *SPON1* |
| *CSMD1* | *SQSTM1* |
| *CYCS* | *ST18* |
| *DLC1* | *STK32B* |
| *DMXL1* | *TBXAS1* |
| *ECHDC3* | *TGM6* |
| *FBXL7* | *THSD4* |
| *FERMT2* | *TMCO4* |
| *GAB2* | *TREM2* |
| *GTF2H3* | *TREML2* |
| *HDAC9* | *TSPAN13* |
| *HECW1* | *UGT1A10* |
| *HS3ST1* | *UGT1A8* |
| *INPP5D* | *USP6NL* |
| *IQGAP2* | *ZCWPW1* |
| *LUZP2* | |
| *MPZL1* | |
| *MS4A4A* | |
| *MS4A4E* | |
| *MS4A6A* | |
